# Candidate undecaprenyl phosphate translocases enable conditional microbial fitness and pathogenesis

**DOI:** 10.1101/2022.02.04.479082

**Authors:** Brandon Sit, Veerasak Srisuknimit, Karthik Hullahalli, Emilio Bueno, Felipe Cava, Matthew K. Waldor

**Affiliations:** Department of Microbiology, Harvard Medical School; Boston, USA; Division of Infectious Diseases, Brigham and Women’s Hospital; Boston, USA; Department of Molecular Biology, Umea University; Umea, Sweden; Department of Immunology and Infectious Diseases, Harvard T. H. Chan School of Public Health; Boston, USA; Howard Hughes Medical Institute; Bethesda, USA

**Author notes:** Department of Biology, Massachusetts Institute of Technology; Cambridge, USA. These authors contributed equally to this work (Brandon Sit, Veerasak Srisuknimit).

## Abstract

The mechanisms that enable adaptation of peptidoglycan, the structural unit of the bacterial cell wall, to shifting extracellular conditions such as pH remain largely unknown. Here, we identify a DUF368-containing membrane protein in the cholera pathogen *Vibrio cholerae* that is critical for pathogenesis and alkaline fitness. *V. cholerae* and *Staphylococcus aureus* lacking their cognate DUF368-containing protein have pH-dependent cell wall defects consistent with surface accumulation of undecaprenyl phosphate (C55-P), an essential lipid carrier for the biogenesis of peptidoglycan and other key bacterial cell surface polymers. In both species, DUF368-containing proteins exhibit synthetic genetic interactions with putative transporters from the DedA family, suggesting these proteins represent complementary long-sought C55-P translocases that enable envelope maintenance functions critical for microbial fitness within and outside the host.

**One-Sentence Summary:** DUF368-containing and DedA-family proteins are undecaprenyl phosphate transporter candidates and are required for bacterial alkaline fitness and pathogenesis.

## Main Text

Microbes must frequently adapt to shifting environmental conditions, including pH. One feature of the bacterial response to pH is reorganization and altered biosynthesis of peptidoglycan (PG), the muropeptide polymer located outside the bacterial membrane that constitutes the bacterial cell wall and maintains cell shape integrity (*1–3*). However, the enzymes and transporters underlying these processes, especially at alkaline pH, are not well-defined. Even in neutral conditions, some gene product(s) required for transmembrane transactions critical for PG biogenesis remain unknown. One such step is the recycling of undecaprenyl pyrophosphate (C55-PP), the lipid carrier required for the transmembrane transport of the sugar-linked pentapeptide subunits of PG (*4*) (Fig. 1A). C55-PP is the membrane-embedded component of the PG precursor Lipid II, which is moved across the inner membrane by the transporter MurJ (*5*). After Lipid II flipping and detachment of the muropeptide, C55-PP recycling is initiated by its hydrolysis to undecaprenyl phosphate (C55-P) by membrane-associated proteins including UppP/BacA and the PAP2-domain proteins PgpB, YbjG, and LpxT (*4*). Then, C55-P is presumably flipped back into the cytosol to complete recycling. C55-P recycling is not only a key step in PG biosynthesis, but required for the assembly of a diverse set of cell surface glycopolymers including wall teichoic acids (WTA), certain LPS modifications, and capsules (*4*). While UppP/BacA has been proposed to also function as a C55-P translocase (*6, 7*), the protein(s) responsible for C55-P internalization have not yet been identified.

**Figure 1.**
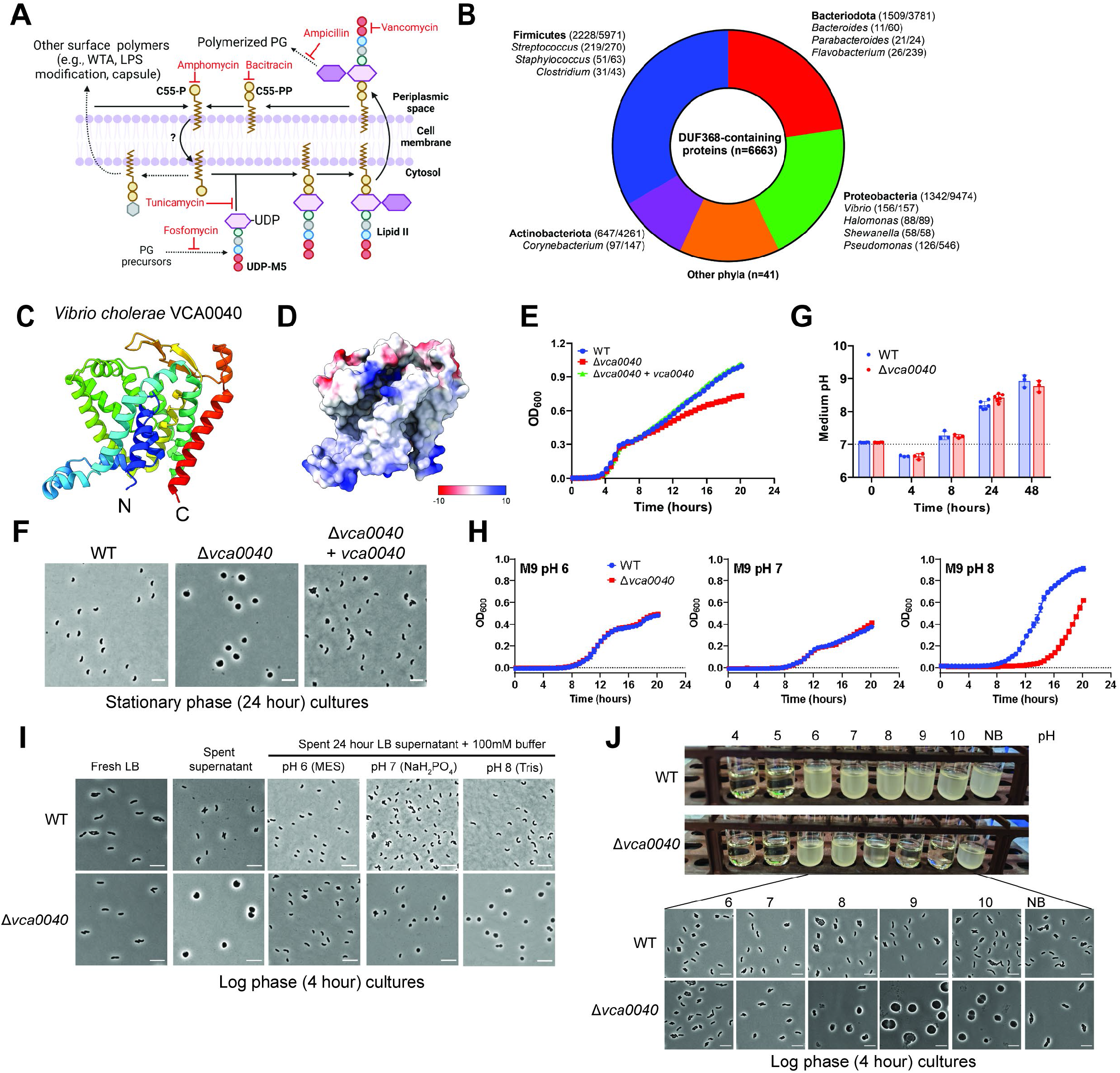
VCA0040 is required for *V. cholerae* cell shape maintenance at alkaline pH. (A): C55-P usage and recycling in bacteria. PG intermediates and sites of antibiotic action are labeled in black and red, respectively. Dashed arrows represent multiple enzymatic and/or transport steps. Grey hexagon represents a range of possible sugars linked to C55-P for downstream glycopolymer assembly. (B) Conservation of DUF368 (PF04018) in bacterial phyla and selected genera. Colored segments and associated labels denote phyla with substantial DUF368 conservation. Numbers next to phylum/genus names indicate the number of sequenced unique clade genomes with a DUF368 protein. (C and D) Predicted ribbon (C) and electrostatic surface colored (D) structures of VCA0040. (E and F) Growth (E) and morphology (F) of Δ*vca0040 V.* cholerae in LB. (G) pH of *V. cholerae* grown in LB medium. (H) *V. cholerae* growth in M9 medium buffered to the indicated pH. (I) Effects of buffered spent supernatant on *V. cholerae* morphology in log phase. (J) Effects of pH on *V. cholerae* growth (top) and morphology (bottom) during log phase in LB medium buffered with 50 mM Na_2_HPO_4_. NB, non-buffered. Scale bars, 5 μm.

*Vibrio cholerae*, the Gram-negative causal agent of the diarrheal disease cholera, is exposed to dramatic changes in pH and ionic context as it enters, transits, and exits the host gastrointestinal tract. A variety of sensing and signaling networks allow *V. cholerae* to adapt to changing environments. For example, the pathogen responds to altered salinity and pH by using a sodium motive force (SMF) instead of a proton motive force (PMF) to power protein secretion and flagellar-dependent motility, and regulate virulence gene expression (*8*). *V. cholerae*’s PG composition is thought to be influenced by environmental inputs, but the effects of pH on *V. cholerae* cell wall biology are unknown (*9*).

A recent *in vivo* transposon-insertion sequencing screen for determinants of intestinal colonization in a contemporary *V. cholerae* clinical isolate identified numerous genes not previously linked to *V. cholerae* pathogenesis, including several loci of unknown function (fig. S1A) (*10*). One of these genes, *vca0040* (*N900_RS14215*), was selected for further study because similar datasets (*11–13*) suggested a universal requirement for this gene in *V. cholerae* intestinal colonization. VCA0040 is predicted to be a multi-pass inner membrane protein, containing a domain of unknown function (DUF368, PF04018) that is conserved across the *Vibrionaceae* and thousands of additional Gram-negative, -positive, and archaeal species that occupy a wide range of niches (Fig. 1B, fig. S1B-E, table S1 and S2). Structural modeling of VCA0040 revealed a large putative cleft in the protein, characteristic of domains with ligand binding and/or transport activity (Fig. 1C, D, fig. S2). A *V. cholerae* strain lacking VCA0040 (Δ*vca0040*) exhibited a stationary phase growth defect (Fig. 1E, fig. S3A). In stationary phase LB cultures, Δ*vca0040* cells exhibited a striking cell shape defect, becoming large spheres (Fig. 1F, fig. S3B, C). These sphere-shaped *V. cholerae* were viable and gave rise to normal rod-shaped daughter cells (fig. S3D).

We reasoned that a stationary phase-specific factor could be triggering the shape defect in Δ*vca0040* cells. Indeed, exposure of exponentially-growing Δ*vca0040* cells to cell-free spent supernatants from stationary phase cultures rapidly induced sphere formation (fig. S3E). Heat-labile and high molecular weight factors, as well as D-amino acids, which modulate stationary phase PG composition (*14*), were excluded as candidates (fig. S3E, F). In LB cultures, *V. cholerae* entry into stationary phase is accompanied by media alkalinization (Fig. 1G). Since only minimal shape defects were observed in neutral buffered M9 medium (fig. S3B), we hypothesized Δ*vca0040* cells were alkaline-sensitive. Buffering M9 samples revealed a growth defect at pH 8, but not pH 6 or 7, and buffering the cell-free spent supernatant to pH 6 or 7 ablated the sphere-induction phenotype (Fig. 1H, I). In buffered LB, Δ*vca0040* cells exhibited growth and cell shape defects at pH ≥ 8, but not at pH ≤ 7 (Fig. 1J). Thus, VCA0040 is required for *V. cholerae* cell shape integrity in alkaline conditions.

Since PG is required for bacterial cell shape maintenance, we measured the amount and composition of the PG in Δ*vca0040* cells. The Δ*vca0040* mutant had 1.5-2x less PG than the wild-type (WT) and modest crosslinking defects with concurrent accumulation of the PG precursor UDP-N-acetylmuramyl pentapeptide (UDP-M5) (Fig. 2A, B, fig. S4A-C) suggestive of a PG biosynthesis defect upstream of crosslinking. Δ*vca0040*’s PG defects were present at neutral pH and exacerbated by exposure to alkaline conditions. During sphere formation in Δ*vca0040 V. cholerae,* there was a ∼10-fold induction of *pgpB*, which encodes a C55-PP phosphatase, suggesting that the absence of VCA0040 impacts PG recycling (fig. S5, table S5). Despite these signals of PG disruption, the Δ*vca0040* mutant only had minor elevations (≤2x) in minimal inhibitory concentrations (MICs) (table S3) to PG-targeted antibiotics, likely because these molecules do not inhibit the step in PG biosynthesis impacted by VCA0040.

**Figure 2.**
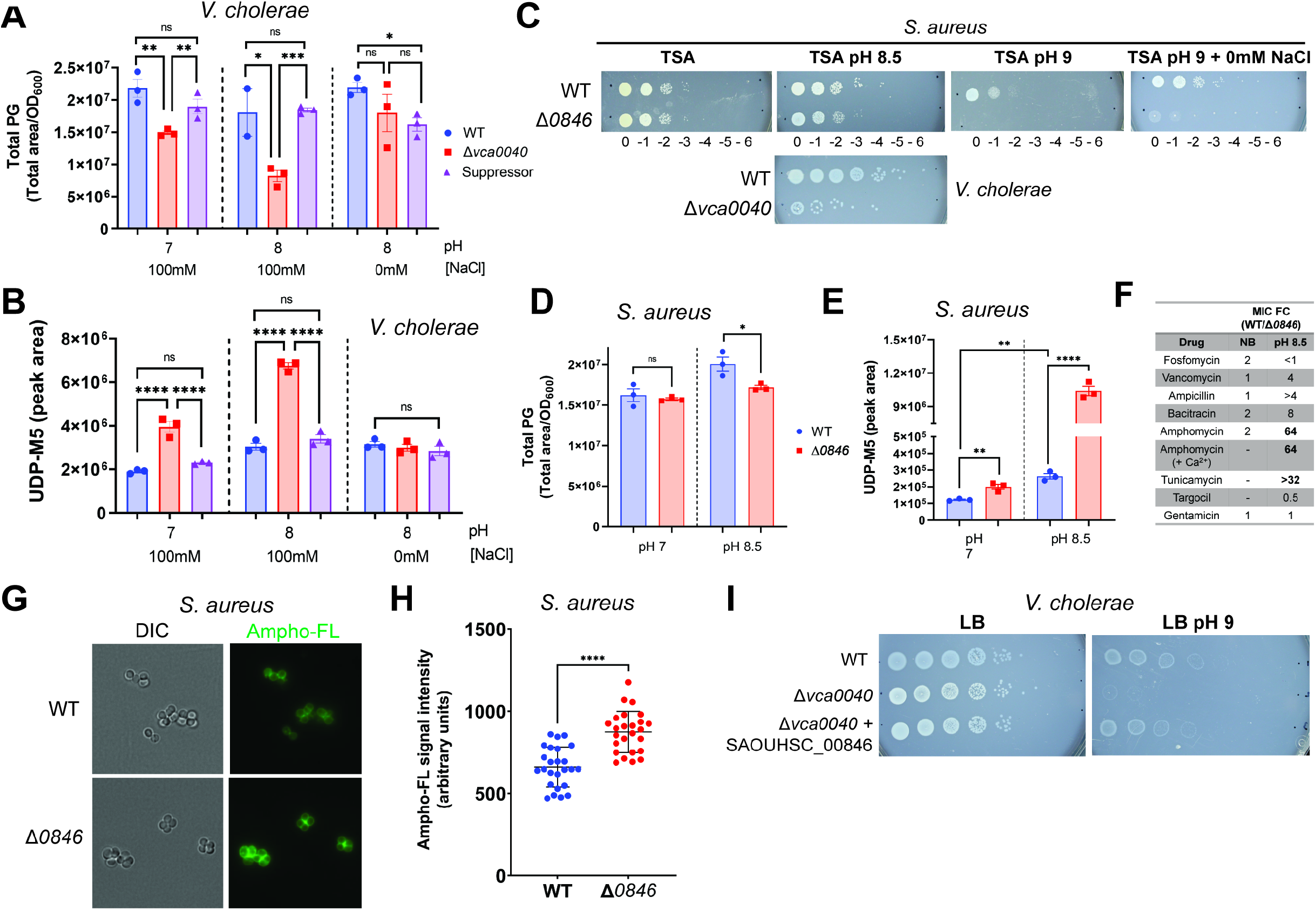
Bacteria lacking DUF368 proteins have reduced PG, increased UDP-M5 and sensitivity to amphomycin. (A and B) Total PG (A) and intracellular UDP-M5 (B) in *V. cholerae* grown in M63 minimal medium at the indicated pH. Suppressor: Δ*vca0040/ΔsecDF1.* (C) Alkaline growth of Δ*SAOUHSC_00846 S. aureus*. (D and E) Total PG (D) and intracellular UDP-M5 (E) in *S. aureus* grown in TSB + 100 mM bicine at the indicated pH. (F): MIC fold change (FC) of Δ*SAOUHSC_00846 S. aureus*; NB, non-buffered, -, not performed. (G): Staining of *S. aureus* with fluorescein-conjugated amphomycin (Ampho-FL). (H): Quantification of ampho-FL signal. (I): Growth of Δ*vca0040 V. cholerae* expressing *SAOUHSC_00846* from the *vca0040* chromosomal locus. *p<0.05, **p<0.001, ***p<0.0005, ***p<0.00005 (unpaired student’s two-tailed t-test). ns, not significant. Where necessary, Δ*SAOUHSC_00846* is abbreviated to Δ*0846*.

To leverage PG-specific antibiotics that cannot be used in Gram-negative bacteria due to the outer membrane, we focused our study on the Gram-positive *Staphylococcus aureus* DUF368-containing protein (SAOUSHC_00846), which has a similar predicted structure to VCA0040 (Fig. 1A, fig. S2). Akin to Δ*vca0040 V. cholerae, S. aureus* lacking *SAOUSHC_00846* had an alkaline growth defect, reduced PG quantity, and accumulation of UDP-M5 as well as decreased cross-linkage (Fig. 2C-E, fig. S4D, E). In alkaline, but not neutral conditions, the Δ*SAOUSHC_00846* mutant was far more sensitive (>64x) to amphomycin than the WT, whereas its sensitivity to most other cell-wall targeting antibiotics only increased modestly (Fig. 2F, table S4). Amphomycin is a lipopeptide antibiotic that specifically binds C55-P and inhibits its recycling (Fig. 1A) (*15*), and is known to induce UDP-M5 accumulation and decrease PG crosslinking in *S. aureus* (*16*). Δ*SAOUHSC_00846* was also more sensitive to tunicamycin, which inhibits the first committed steps of WTA and PG synthesis, both of which are C55-P-dependent (Fig. 1A, 2F, table S4) (*17*). The antibiotic and PG phenotypes are consistent with accumulation of surface C55-P in the outer leaflet of the cell membrane in the Δ*SAOUHSC_00846* mutant. To directly visualize surface-exposed C55-P, we synthesized fluorescein-conjugated amphomycin (ampho-FL) and used it to label live bacterial cells (*18*). Ampho-FL-labeled Δ*SAOUHSC_00846 S. aureus* exhibited increased signal relative to WT bacteria, suggesting that C55-P is not efficiently internalized in the mutant (Fig. 2G, H). Heterologous expression of *SAOUHSC_00846* in Δ*vca0040 V. cholerae* rescued the alkaline defect (Fig. 2I, fig. S4F), indicating that DUF368 protein function is conserved. The phenotypes of DUF368-deficient *S. aureus* and *V. cholerae* strongly suggest DUF368 domains are involved in C55-P recycling, likely as C55-P transporters.

Analyses of spontaneous suppressors of Δ*vca0040 V. cholerae*, which have distinctive colony morphology (fig. S6A) revealed that VCA0040’s requirement for *V. cholerae* fitness is controlled by [Na^+^] in addition to pH. Suppressor colonies lacked stationary phase cell shape defects and whole-genome sequencing revealed four suppressor genes, three of which (*secD1, secF1,* and *ppiD*) function in Sec-mediated protein secretion (fig. S6B). The fourth, *yfgO,* is a protein of unknown function from the AI-2E transporter family, members of which have recently been implicated in PG biosynthesis and Na^+^/H^+^ antiport (*19, 20*). Deletions of each gene rescued Δ*vca0040*’s shape defects, even though stationary culture pH was alkaline in all suppressors (fig. S6C, D). In *V. cholerae*, SecDF is duplicated as SecD1/F1 and SecD2/F2, ancillary complexes that differentially couple SecYEG activity to the SMF or PMF, respectively (fig. S6E) (*21*). The activity of at least one SecDF pair was required for *V. cholerae* viability and deletion of *secD2/F2* did not ameliorate Δ*vca0040’s* stationary phase cell shape defect (fig. S6C, F, G). We initially hypothesized that altered Sec substrate specificity could explain the identification of *secD1/secF1* as suppressors. However, the Δ*secDF1 V. cholerae* proteome was unchanged from that of WT cells apart from expected increases in SecDF2, suggesting the suppressive effects of *secD1/secF1* loss were attributable to reduced Na^+^ flux (fig. S6H, table S6). Indeed, Na^+^-free, but not K^+^-free conditions suppressed the alkaline growth and PG defects of Δ*vca0040 V. cholerae* (Figs. 2A, B, 3A, figs. S5C, S6I, J). Thus, alkaline pH and Na^+^ both contribute to the environmental control of the defect of DUF368-deficient *V. cholerae*. The Na^+^ requirement appears to be species-dependent, as Δ*SAOUHSC_00846 S. aureus*, although alkaline-sensitive, were not similarly rescued by Na^+^ depletion (Fig. 2C).

**Figure 3.**
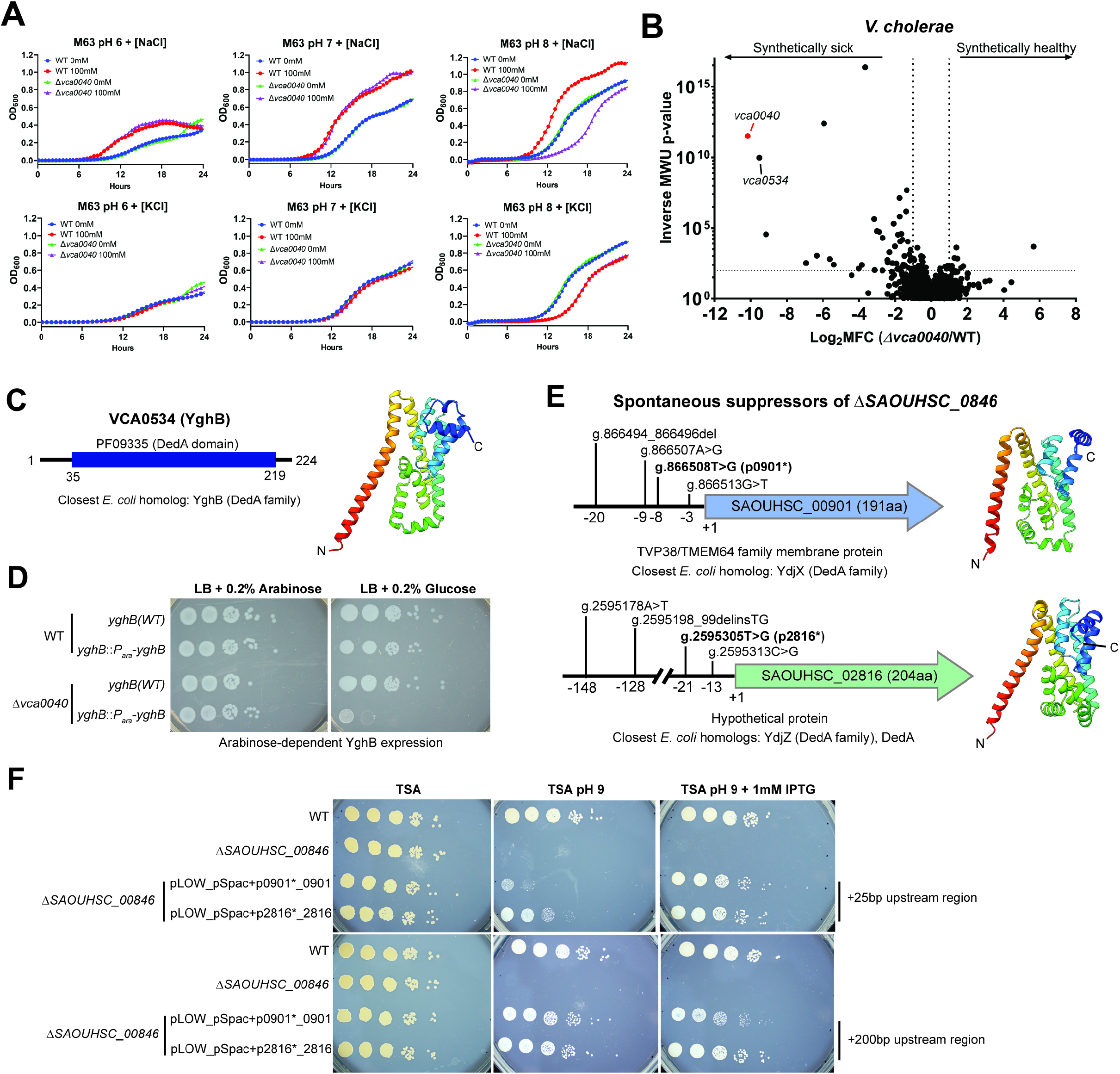
Environmental and genetic modifiers of DUF368 essentiality. (A): Rescue of Δ*vca0040 V. cholerae* in Na^+^-free (top) but not K^+^-free (bottom) alkaline M63 media. (B): Synthetic transposon screen in Δ*vca0040 V. cholerae*, showing log_2_ mean read fold changes (MFC, threshold ±2) and inverse Mann-Whitney U (MWU, threshold 100) test p-values. (C): Domain content and predicted structure of *V. cholerae* YghB. (D): Validation of synthetic lethal *vca0040/yghB* interaction. (E): Spontaneous suppressors of Δ*SAOUHSC_00846* map to two *S. aureus* DedA-family proteins, with predicted structures (right) and specific promoter mutations selected for validation (bold). (F): Rescue of Δ*SAOUHSC_00846* by expression of SAOUHSC_00901 or SAOUHSC_02816 from IPTG-inducible vector (pSpac) and isolated mutant promoters (marked with *).

We next carried out a synthetic transposon screen to define *vca0040*’s genetic network. Many of the identified synthetic sick/lethal interactions were related to cell envelope homeostasis (Fig. 3B, table S7). *vca0040’s* most striking interaction was with *N900_RS16280* (*vca0534,* Supplementary Text), encoding a homolog of the *E. coli* DedA family member YghB (Fig. 3C). Two other DedA proteins are present in *V. cholerae* but were not hits. DedA family transmembrane proteins (PF09335, SNARE-associated Golgi domain) are conserved in all three domains of life, generally required for cell envelope homeostasis, and are suspected to mediate PMF-dependent transport, but their specific substrates are unknown (*22, 23*). VCA0040’s synthetic lethality with YghB was validated by genetic depletion and reciprocal synthetic transposon screening (Fig. 3D, fig. S7A, table S8), and overexpression of YghB rescued the alkaline defect of Δ*vca0040 V. cholerae* (fig. S7B). Strikingly, interactions between DUF368 and DedA proteins were also observed in *S. aureus.* In alkaline-enriched spontaneous suppressors of *S. aureus* Δ*SAOUHSC_00846,* we found frequent promoter region (and no coding region) mutations in two DedA proteins (SAOUHSC_00901 and SAOUHSC_02816) (Fig. 3E). Incorporating these mutations into an IPTG-inducible system rescued *SAOUHSC_00846* even without induction, suggesting the isolated mutations increase expression of DedA-family proteins and compensate for the loss of *SAOUHSC_00846,* as in *V. cholerae* (Fig. 3F). The conserved synthetic interaction between DUF368 and DedA proteins suggests that DedA family members are also required for C55-P transport, consistent with recent reports that eukaryotic DedA proteins may bind lipids, and bacterial DedA mutants show decreases in LPS modifications that require C55 carriers (*23–25*). *V. cholerae* lacking both *yghB* and *vca0040* were viable at acidic pH, suggesting at least one other *V. cholerae* protein can carry out C55-P translocation (fig. S7C). DedA proteins may be more active in different conditions than DUF368 proteins, since opposite from the Δ*vca0040* mutant, a Δ*yghB* mutant was not able to grow without Na^+^, suggesting an SMF-independent (i.e., PMF-dependent) function (fig. S7D). Consistent with the idea of conditional C55-P translocase activity, the outer membrane-permeable *lptD4213 E. coli* strain (*26*), which has only DedA paralogs and not DUF368, had heightened sensitivity to amphomycin and tunicamycin at alkaline pH (table S4).

We used an infant rabbit model, which mimics severe human cholera (*27*), to investigate VCA0040’s role in pathogenesis. Typically, alkaline inocula are used in this model to buffer the acidic stomach environment; to exclude inoculum effects, we carried out mixed infections with 1:1 WT: Δ*vca0040 V. cholerae* inocula at pH 7 or pH 9 (Fig. 4A). Regardless of inoculum pH, the *Δvca0040* mutant had a profound competitive defect (∼100-1000x) versus the WT strain (Fig. 4B). Inoculum pH did not affect total colonization or accumulation of diarrhea-like cecal fluid (CF), the primary marker of disease in this model (fig. S8A, B). In single infections with fluorescent WT or Δ*vca0040 V. cholerae,* the mutant had severe colonization defects (∼1000-fold) in all intestinal segments (Fig. 4C, D). These values overestimate the true colonization of the mutant because animals from each group were co-housed to exclude litter effects, enabling substantial transmission from WT-infected to Δ*vca0040*-infected animals (fig. S8C). Transmission likely explains why CF accumulation in Δ*vca0040*-infected animals was lower than WT but did not reach statistical significance (Fig. 4E). Measurements of CF from both mixed and singly-infected animals revealed an alkaline pH of 8.5-9 (Fig. 4F). Imaging of CF samples from Δ*vca0040*-infected rabbits revealed detectable large fluorescent spherical cells similar to those observed in alkaline cultures of the Δ*vca0040* mutant (fig. S8D). To corroborate this idea, freshly grown rod-shaped WT or Δ*vca0040 V. cholerae* were incubated in cell-free CF samples. Δ*vca0040,* but not WT, cells became spherical when exposed to CF, suggesting that VCA0040 function is critical for maintenance of *V. cholerae* shape in the *in vivo* milieu (Fig. 4G). An infection defect was not detected for Δ*SAOUHSC_00846 S. aureus* in a murine IV infection/organ abscess model, where the pathogen is unlikely to encounter alkaline environments (fig. S9).

**Figure 4.**
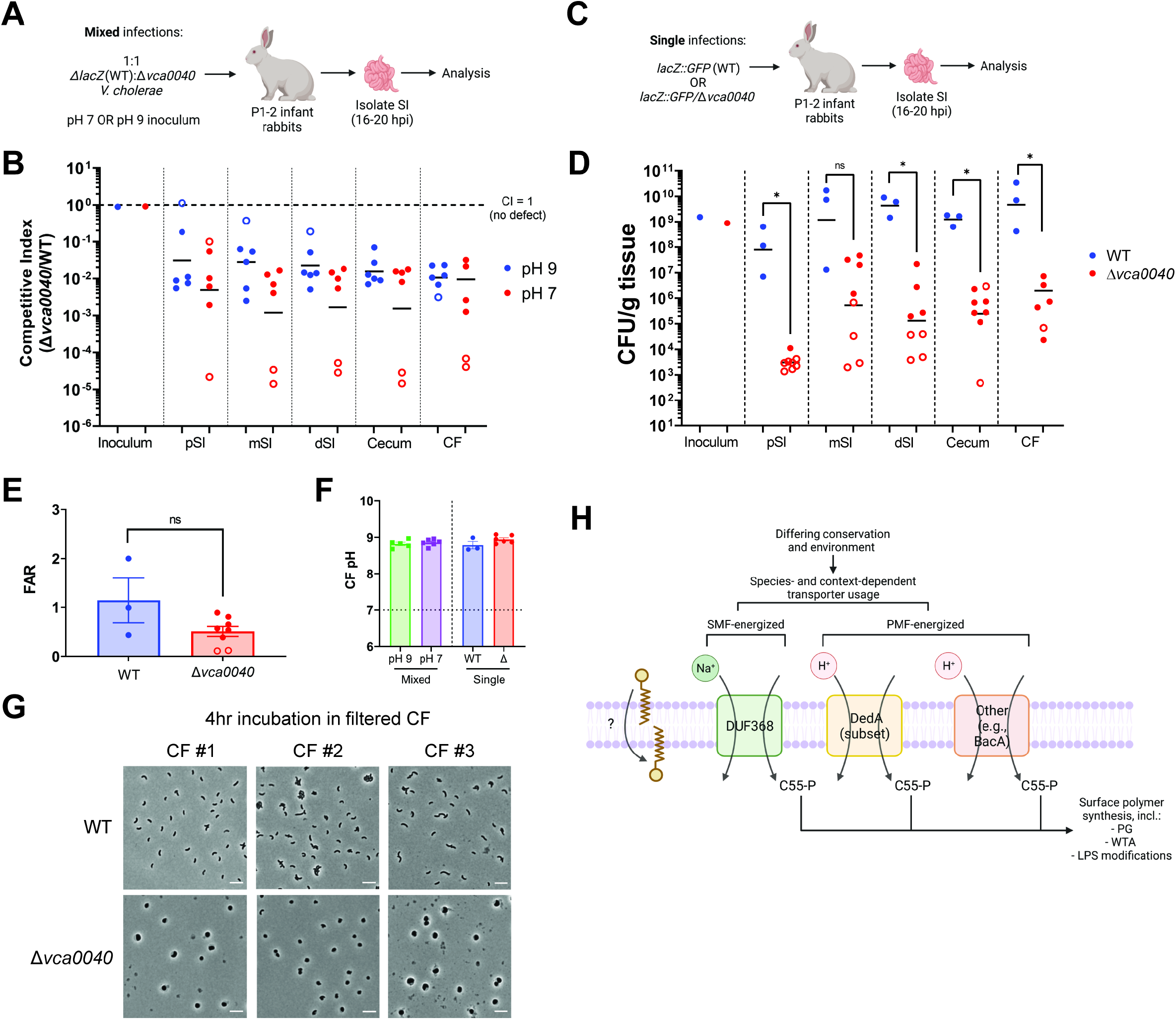
VCA0040 is required for *V. cholerae* pathogenesis. (A and B): Schematic (A), and intestinal competitive indices (B) in mixed infections. (C-E): Schematic (C), intestinal colony-forming units (D), and fluid accumulation ratio (FAR) (E) in single infections. (F): Cecal fluid (CF) pH from infected animals. (G): Incubation of lab-grown *V. cholerae* with filtered CF samples from three different rabbits. SI: small intestine, pSI: proximal SI, mSI: medial SI, dSI: distal SI. Open circles indicate animals with limit of detection measurements. *p<0.05 (Mann-Whitney U test). ns, not significant. (H): Proposed model for microbial C55-P translocation and its dependence on species and environment. In *V. cholerae*, VCA0040 (DUF368) and YghB (DedA) are likely the primary translocases due to their synthetic lethality. VCA0040 is likely SMF-dependent, as Δ*yghB* cannot grow with Na^+^. YghB is likely PMF-dependent, because Δ*vca0040* is alkaline (low PMF)-sensitive. The acidic (high PMF) rescue of a *yghB/vca0040* double mutant suggests that additional PMF-dependent translocases may (e.g., BacA/UppP) exist.

We propose that DUF368-containing and DedA family proteins are C55-P translocases that provide functional redundancy in this crucial step of envelope maintenance. The apparent environmental specificity of DUF368-containing and DedA proteins (and potentially other proteins such as BacA/UppP) could support C55-P flux in diverse microbial niches (Fig. 4H). Environmental tuning of C55-P recycling through C55-P translocase regulation may occur via bioenergetic diversification, as our data suggest that *V. cholerae* VCA0040 is SMF-controlled and YghB is PMF-controlled. The apparent enrichment of DUF368 in known halophiles such as *V. cholerae, S. aureus,* and much of the archaeal class Halobacteria suggests DUF368 may facilitate microbial adaptation to high-salt environments (table S1). Alongside the complex pattern of conservation of redundant translocases (fig. S10, table S9, Supplementary Text), additional inputs and adaptation mechanisms likely also contribute to control of C55-P recycling. For example, although *vca0040* is only required in alkaline conditions in *V. cholerae*, this gene is essential in the related pathogen *V. parahaemolyticus* even at neutral pH, despite conservation of YghB and suppressor loci (fig. S11) (*28*). Although DUF368-containing proteins, renamed PopT for polyprenol phosphate transporter in a concurrently submitted study (Roney and Rudner, *bioRxiv* 2022), are restricted to bacteria and archaea, DedA family members are widely present in eukaryotes, including humans. Thus, our findings may impact understanding of translocation of polyprenol phosphates (e.g., dolichol phosphate) across the kingdoms of life.

## Acknowledgements

We thank members of the Waldor lab for helpful comments and discussions during the study. We also thank the Thermo Fisher Scientific Center for Multiplexed Proteomics at Harvard Medical School (http://tcmp.hms.edu), especially Dr. Rachel Rodrigues, for assistance with proteomics, and the Microbial Genome Sequencing Center (MiGS) for assistance with RNAseq and whole-genome sequencing. Schematics and model figures were generated with Biorender (Biorender.com). This article is subject to HHMI’s Open Access to Publications policy. HHMI lab heads have previously granted a nonexclusive CC BY 4.0 license to the public and a sublicensable license to HHMI in their research articles. Pursuant to those licenses, the author-accepted manuscript of this article can be made freely available under a CC BY 4.0 license immediately upon publication.

## Funding

Swedish Research Council (FC)

Knut och Alice Wallenbergs Stiftelse (FC)

The Laboratory of Molecular Infection Medicine Sweden (FC)

The Kempe Foundation (FC)

National Institutes of Health RO1AI-042347 (MKW)

Howard Hughes Medical Institute (MKW)

Wellcome Trust (MKW)

Natural Sciences and Engineering Research Council of Canada PGSD3-487259-2016 (BS)

National Institutes of Health 1F31AI156949-01 (KH)

## Author Contributions

Conceptualization: BS, MKW, VS

Methodology: BS, VS, KH, EB, FC, MKW

Investigation: BS, VS, KH, EB

Visualization: BS, VS

Funding acquisition: FC, MKW

Project administration: MKW

Supervision: FC, MKW

Writing – original draft: BS, VS, MKW

Writing – review & editing: BS, VS, KH, EB, FC, MKW

All data are available in the main text or the supplementary materials.

## Supplementary Materials for

**This PDF file includes:**

Materials and Methods
Supplementary Text
References (29–47)
Figs. S1 to S11
Tables S1 to S11

### Materials and Methods

Note: Parts of this section are adapted from the thesis of B.S (*29*).

#### Bacterial strains, media and growth conditions

All *V. cholerae* strains used in this study are derivatives of HaitiWT, a spontaneous streptomycin-resistant variant of a clinical isolate from the Haiti cholera epidemic (*30*). All *S. aureus* strains used in this study are derivatives of HG003 (itself a derivative of NCTC8325). Strain and plasmid information is listed in Table S10. Genome annotation of HaitiWT and a batch BLAST dictionary assigning putative VC locus tags (the conventional *V. cholerae* gene naming system) to HaitiWT loci is given in Table S11. All constructed strains were verified by Sanger sequencing of the targeted locus (Genewiz, USA).

The following media were used in this study: Lysogeny broth (LB) Miller (10 g/L NaCl) (BD Biosciences, USA), M9 minimal media (12.8 g/L Na_2_HPO_4_·7H_2_O, 3 g/L KH_2_PO_4_, 0.5 g/L NaCl, 1 g/L NH_4_Cl, 1 mM MgSO_4_ and 10 µM CaCl_2_) (lab-made), M63 minimal media (2 g/L (NH_4_)_2_SO_4_, 13.6 g/L KH_2_PO_4_, 0.5 mg/L FeSO_4_·7H_2_O and 1 mM MgSO_4_) (US Biologicals, USA), and tryptic soy broth and agar (TSB/TSA) (BD Biosciences, USA). M9 was supplemented with 0.4% glucose and pH-adjusted with NaOH. M63 was supplemented with 2% glucose and pH-adjusted with KOH. LB was buffered with 25 mM Na_2_HPO_4_ or 100 mM Tris and pH-adjusted with NaOH or HCl. TSB was buffered with 100 mM bicine. Plates were used at 1.5% final agar concentration. Where necessary, the following antibiotics or supplements were used: streptomycin (Sm, 200 μg/mL), carbenicillin (Cb, 50 µg/mL), kanamycin (Kn, 50 µg/mL), neomycin (50 ug/mL), targocil (10 μg/mL), diaminopimelic acid (DAP, 0.3 mM), 5-bromo-4-chloro-3-indolyl β-d-galactopyranoside (X-gal, 60 µg/mL), arabinose (0.2%), isopropyl ß-D-1-thiogalactopyranoside (IPTG, 1 mM).

For routine culture, *V. cholerae* were grown in non-buffered LB at 37°C and *S. aureus* were grown in non-buffered TSB at 37°C. Cultures for spent supernatant analysis in *V. cholerae* were obtained by subculturing 37°C-grown overnight cultures 1:1000 in fresh media and growing for 24 hours at 30°C. To collect spent supernatants, cultures were centrifuged for 10 minutes at 5000 rcf at 4°C. Supernatants were transferred to a new tube, re-centrifuged, and sterile filtered with 0.22 µm syringe filters into new tubes. Supernatants were stored at 4°C for future use. Cultures used for pH quantification were centrifuged for 10 minutes at 5000 rcf before measurement. All pH measurements, including for media and buffer formulation, were performed with a pH meter (Thermo ORION, USA) freshly calibrated to two appropriate pH standards from 4, 7, and 10.

#### Bioinformatic methods

Information on the loci, genomes, and accession numbers for genes and proteins from this study is available in Table S10.

#### Phylogenetic analysis

To identify sequences with annotated DUF368 domains, Annotree (*31*) was used with the Pfam identifier PF04018 at a cutoff e-value of 1×10^-30^. The results from this search are included in Table S1. Phyla with “_X” names were manually collated into a single phylum for ease of analysis. In a related effort, to identify precise homologues of *vca0040*, we used a HMMER search with the VCA0040 sequence from WT *V. cholerae* (Table S2). To analyze the relative distributions of PF04018, PF09335 (DedA), and PF02673 (BacA), we used Annotree at a cutoff e-value of 1×10^-15^. Differential species lists with all possible combinations of conservation were generated with a custom Venn diagram generator (https://bioinformatics.psb.ugent.be/webtools/Venn/) and are provided in Table S9.

#### Structural modeling

Structural prediction of DUF368 and DedA-family proteins in this study was performed with AlphaFold2 on the CoLabFold publicly accessible interface (*32*). Sequences were modeled as monomers using mmseqs2 for multiple sequence alignment. Structures were ranked by pLDDT and the top-ranked structures were visualized with ChimeraX (UCSF, USA).

#### Cloning, vectors, and strain construction

##### V. cholerae

Cloning of expression vectors and deletion plasmids was performed by standard isothermal assembly techniques with 25 bp overhangs on each fragment (HiFi DNA Assembly Kit, NEB, USA). *V. cholerae* mutants were generated by allelic exchange as previously described with the suicide vector pCVD442 bearing 500-700 bp upstream and downstream homology arms of the targeted locus (*33*). Either SM10λpir or MFDλpir E. coli (a DAP auxotroph) were used as the donor strain with identical conjugation conditions apart from DAP supplementation. Single crossover transconjugants were isolated by selective plating on LB + Sm/Cb and passaged in 10% sucrose overnight at RT to select for double crossover events. Cells were plated on LB + Sm, re-patched onto LB + Sm and LB + Sm/Cb, and Sm^R^Cb^S^ colonies screened by colony PCR for the successful deletion. For deletions involving *vca0040*, *vca0040* was always deleted last and at least 2 clones per strain were stocked and verified for the correct phenotype to guard against spontaneous suppressor formation. For native *vca0040* expression from the ectopic chromosomal site, the full coding sequence along with 350 bp upstream was cloned to include the native promoter and inserted at a known neutral genomic location in *V. cholerae* (*34*). For expression of *SAOUHSC_00846* from the *vca0040* locus in Δ*vca0040 V. cholerae*, we used allelic exchange to replace the *vca0040* coding sequence with a *SAOUHSC_00846* sequence codon-optimized for *V. cholerae*. For depletion of *yghB* from Δ*vca0040 V. cholerae*, we used pAM299, a suicide vector which replaces the native allele of targeted gene with an arabinose-inducible copy (*35*). For overexpression studies, we used pBAD18 for plasmid-based arabinose induction (*36*).

##### S. aureus

*S. aureus SAOUHSC_00846* knockout was generated with a one-step allelic exchange using pTarKO plasmid bearing 1000 bp upstream and downstream homology arms of the targeted locus flanking a kanamycin cassette as previously described (*37*). Briefly, assembled pTarKO plasmid was electroporated into electrocompetent *S. aureus* RN4220 tarO_off_ and selected for double crossovers on TSB with kanamycin, neomycin and targocil. The kanamycin insertion in *SAOUHSC_00846* was then transduced to *S. aureus* HG003 with phage phi85. For overexpression studies, we used pLOW for plasmid-based IPTG induction (*38*). Assembled plasmids were electroporated into RN4220 WT and transduced to *S. aureus* HG003 Δ*SAOUHSC_00846* with phage phi85.

##### V. parahaemolyticus

*V. parahaemolyticus* RIMD2210633 mutants were constructed with a similar allelic exchange system to *V. cholerae* as previously described, with the suicide vector pDM4 (*28*).

#### Growth assays

##### Growth curves in liquid medium

For automated OD_600_ measurements, 1 mL of a saturated 37°C LB overnight *V. cholerae* culture was washed once with 1 mL fresh media (specific to the experiment) and resuspended in 1mL fresh media. Resuspended bacteria were diluted to a starting dilution of 1:4000 by performing a 1:100 dilution into 1 mL fresh media and a subsequent 1:40 dilution into 195 µL of specific media aliquoted into sterile 96 well plates (Corning, USA). At least 3 technical replicates per strain and media condition were run per plate along with at least 3 blank media wells. Growth curves were performed in a BioTek Epoch2 spectrophotometer with shaking and OD_600_ readings were taken every 10 minutes for 20-24 hours. Data for each condition were averaged across technical and biological replicates corrected against the baseline blank OD_600_ values.

##### Overexpression vector plate dilution assays on solid medium

*V. cholerae* strains were grown overnight at 37°C in LB with addition of 50 µg/mL carbenicillin for strains with pBAD18. The cultures were diluted 1:100 into 5 mL of LB (supplemented with 50 µg/mL carbenicillin and 0.2% arabinose for strains with pBAD18) and grown at 37 °C to an OD_600_ of ∼0.7. Cells were normalized to an OD_600_ of 0.1 and then serially 10-fold diluted six times. 5 µL of the dilution series were plated on LB agar 100 mM Tris pH 9 (with 0.2% arabinose, 0.2% glucose, or no sugar added) and incubated at 30°C for 18-24 hours.

*S. aureus* strains were grown overnight at 37 °C in TSB with addition of 10 µg/mL erythromycin for strains with pLOW. The cultures were diluted 1:100 into 5 mL of TSB (supplemented with 10 µg/mL erythromycin and 1 mM IPTG for strains with pLOW) and grown at 37 °C to an OD_600_ of ∼0.7. Cells were normalized to an OD_600_ of 0.1 and then serially 10-fold diluted six times. 5 µL of the dilution series were plated on TSB agar 100 mM Tris pH 9 (with or without 1 mM IPTG) and incubated at 37°C for 18-24 hours.

#### Live and single timepoint phase-contrast microscopy

For single timepoint imaging of live cells, samples from the indicated cultures were concentrated as necessary and immobilized on 0.8% agarose pads in sterile PBS on glass slides (Gene Frames, Thermo, USA) and dried before coverslip placement. For time-lapse imaging of live cells, samples were spotted on 0.8% agarose pads in sterile LB before imaging in a temperature-controlled chamber. Cells were imaged with a Nikon Eclipse Ti microscope equipped with an Andor NeoZyla camera and a 100× oil phase 3 1.4-numerical-aperture (NA) objective. Image analysis was performed with ImageJ. Images in manuscript figures are representative of at least 10 fields from the same sample (>200 cells) and multiple independent replicate cultures.

#### Sphere formation and incubation assays

To induce sphere formation, *V. cholerae* 37°C overnight cultures (where Δ*vca0040* cells are still largely rod shaped) were back-diluted 1:100 into the indicated supernatant or fresh media and grown for 4 hours shaking at 200 rpm at 30°C. Then, cells were imaged as described above. For the D/L-Ala treatment assay, overnight cultures were expanded in fresh LB for 90 minutes prior to spike-in of the amino acid for 1 hour.

#### Minimal inhibitory concentration assays

##### V. cholerae

To quantify MICs for various antimicrobial agents, 37°C overnight cultures of *V. cholerae* were diluted 1:100,000 in fresh media. Diluted cultures were used to inoculate 96-well plates containing twelve 2-fold dilutions of the indicated agent in LB medium at a ratio of 50 µL culture: 50 µL medium. Four technical replicates were performed per strain per dilution. Plates were incubated for 24 hours at 37°C and MIC values were read as the first dilution where no turbidity was observed. For repeat assays, MICs were performed with independent overnight cultures.

##### S. aureus and E. coli

37°C overnight cultures of *S. aureus* or *E. coli* were diluted to OD_600_ = 0.01 and then further diluted 1:100 in fresh TSB media. Diluted cultures were added to 96-well plates containing 2-fold dilutions of the indicated agent in TSB medium or TSB medium 5 mM Tris pH 8.5 at a ratio of 75 µL culture: 75 µL media. Plates were incubated for 24 hours at 30°C (37°C for *E. coli*) with shaking. MIC values were read off as the first dilution where no turbidity was observed.

#### Peptidoglycan characterization

##### Preparation of *V. cholerae* and *S. aureus* sacculi

For *V. cholerae,* strains were grown overnight at 37°C in LB + 200 µg/mL streptomycin. For each sample, 500 µL of overnight culture were collected and centrifuged at 5000 rcf for 5 min, washed once with 500 µL of the corresponding media, and resuspended in 500 µL of the corresponding media. This culture was then added to 50 mL of M63 media (pH 7 with 100 mM NaCl, pH 8 with 100 mM NaCl or pH 8 with 0 mM NaCl) supplemented with 2% glucose and 200 µg/mL streptomycin and incubated at 30°C for approximately 6 hours until the OD_600_ reached ∼0.5. Bacteria cells were collected by centrifugation at 6000 rcf for 10 minutes at 4°C and resuspended in 1.5 mL of 1 x PBS. The resuspension was added dropwise into 1.5 mL of boiling 5% SDS solution. The mixture was boiled for 1 hour and stirred for 2 more hours after the heat was turned off.

For *S. aureus,* strains were grown overnight at 37°C in 3 mL Tryptic Soy Broth (TSB). 500 µL of the overnight culture was added to 100 mL of TSB without pH adjustment or TSB + 100 mM bicine pH 8.5 and incubated at 37°C for ∼2 hours until the OD_600_ reached 0.5. The bacterial cells were collected by centrifugation at 5000 rcf for 10 min at 4°C and resuspended in 1.5 mL of 1 x PBS. Resuspended samples were boiled as for *V. cholerae*.

##### Total PG and crosslinking quantification from sacculi samples

Peptidoglycan was extracted from boiled samples as described previously for Gram-negative and Gram-positive organisms (*39, 40*). Once boiled, cell wall material was pelleted by ultracentrifugation and washed with water. Clean sacculi were digested with muramidase (100 μg/ml) and soluble muropeptides reduced using 0.5 M sodium borate pH 9.5 and 10 mg/mL sodium borohydride. The pH of the samples was then adjusted to 3.5 with phosphoric acid. UPLC analyses were performed on a Waters-UPLC system equipped with an ACQUITY UPLC BEH C18 Column, 130Å, 1.7 μm, 2.1 mm × 150 mm (Waters Corporation, USA) and identified at Abs. 204 nm. Muropeptides were separated using a linear gradient from buffer A (phosphate buffer 50 mM pH 4.35) to buffer B (phosphate buffer 50 mM pH 4.95 methanol 15% (v/v)). Identification of individual peaks was assigned by comparison of the retention times and profiles to validated chromatograms. The relative amount of each muropeptide was calculated by dividing the peak area of a muropeptide by the total area of the chromatogram. The abundance of PG (total PG) was assessed by normalizing the total area of the chromatogram to the OD_600_. The degree of cross-linking was calculated as described previously (*41*).

##### Intracellular UDP-M5 quantification

For *V. cholerae,* strains were grown overnight at 37°C in LB + 200 µg/mL streptomycin. For each sample, 500 µL of overnight culture were collected and centrifuged at 5000 rcf for 5 min, washed once with 500 µL of the corresponding media, and resuspended in 500 µL of the corresponding media. 80 µL of this resuspension was then added to 8 mL of M63 media (pH 7 with 100 mM NaCl, pH 8 with 100 mM NaCl or pH 8 with 0 mM NaCl) supplemented with 2% glucose and 200 µg/mL streptomycin and incubated at 30°C for approximately 6 hours until the OD_600_ reached ∼0.5. Bacteria cells were collected by centrifugation at 5000 rcf for 10 minutes at 4°C, washed twice with 1 mL of ice cold 0.9% NaCl, and resuspended in 200 µL of milliQ water. The resuspension was boiled for 30 min, centrifuged at 20,000 rcf for 15 min, and the supernatant was filtered and analyzed. For *S. aureus,* strains were grown overnight at 37°C in 3 mL Tryptic Soy Broth (TSB). 80 µL of the overnight culture was added to 8 mL of TSB without pH adjustment or TSB + 100 mM bicine pH 8.5 and incubated at 37°C for ∼2 hours until the OD_600_ reached ∼0.5. Cultures were then processed as for *V. cholerae*.

Quantification of soluble UDP-M5 muropeptide levels by LC-MS of filtered supernatants was performed as previously described (*42*). Detection and characterization of soluble muropeptides by LC-MS was performed on an UPLC system interfaced with a Xevo G2/XS Q-TOF mass spectrometer (Waters Corporation) using previously reported conditions (*42*). UDP-M5 levels were quantified by integrating peak areas from extracted ion chromatograms (EICs) of the corresponding m/z value.

#### Amphomycin-FL (ampho-FL) synthesis and staining

To synthesize ampho-FL, 5 mg of amphomycin (Cayman Chemical, USA) was dissolved in 200 µL of dimethyl formamide (DMF) and combined with 7 µL of triethylamine. Separately, 3 mg of fluorescein-C5,6-NHS (ThermoFisher, USA) was dissolved in 200 µL of DMF and then added to the amphomycin solution. The reaction mixture was stirred at room temperature in the dark for 24 hours. The solution was diluted in one equal volume of DMSO and purified by reverse-phase HPLC (Agilent 1260 Infinity) using a C18 stationary phase column (Luna 5 µM C18(2) 100 Å, 250 x 10 mm). HPLC conditions were as follows: Phase A: water (0.1% formic acid); Phase B: acetonitrile (0.1% formic acid). Phase B: 0-2 min, 50%; 2-15 min, linear gradient 50%-100%, 15-17 min, 100%. The wavelength of the detector was set at 254 nm. The flow rate was 4.7 mL/min. The mixture of conjugated fluorescein-C5,6 products eluted at 9 min. HPLC eluates were collected in a 50 mL round-bottom flask, concentrated by rotatory evaporation, transferred with DMSO to a tared microcentrifuge tube and lyophilized, resulting in 1.7 mg of ampho-FL (M+1 = 1649.23).

For ampho-FL labeling, overnight cultures of *S. aureus* HG003 WT or Δ*SAOUHSC_0084*6 were diluted 1:100 in fresh TSB media + 100 mM bicine pH 8.5. The new cultures were grown to OD_600_ = 0.5. 1 mL of the culture was centrifuged at 1000 rcf for 5 min. The pellet was resuspended in 500 µL of 1 x Tris-buffered saline (TBS) pH 9. 98 µL of the mixture was transferred to a new tube and added with 2 µL of 10 mg/mL ampho-FL conjugate in DMSO. The mixture was incubated in the dark at room temperature for 10 minutes, washed three times with 500 µL of 1 x TBS pH 9, and resuspended in 100 µL of 1 x TBS pH 9. 10 µL bacteria was then added to a 0.8% agarose pad in TBS pH 9 and imaged as described above. Bacterial cells were imaged using DIC at a 30 ms exposure and ampho-FL signal was captured with the FITC filter at a 1 second exposure. Fluorescence intensity of ampho-FL was quantified using ImageJ. FITC signal profiles were generated by drawing bisecting lines through dividing cells. The FITC signal was recorded at the background to the left of the cells (A), at the left wall peak (B), at the middle walls (C), at the right wall peak (D), and at the background to the right of the cells (E). The average signal was calculated by (B+C+D)/4-(A+E)/2.

#### Transposon-insertion sequencing (TIS)

Generation of transposon libraries in the Δ*vca0040* and Δ*yghB* backgrounds was performed as previously described for HaitiWT *V. cholerae* (*10*). Strains were conjugated to SM10λpir *E. coli* bearing the donor transposon vector pSC189. Due to the moderate conjugation defect of Δ*vca0040*, we used a 5x concentration of the conjugation reactions compared to the other libraries. Reactions were plated on 245 mm^2^ LB+Sm/Kn agar plates to isolate *V. cholerae* transconjugants and incubated overnight at 30°C. Two independent Δ*vca0040* and Δ*yghB* libraries were generated, consisting of approximately 200,000 colonies each and stocked at an OD_600_ ∼10 in LB + 25% glycerol. For synthetic TIS analyses, a frozen aliquot of each library was thawed and used for genomic DNA extraction with the Wizard kit (Promega, USA). DNA libraries were prepared in an identical manner to previous TIS experiments from our group (ref) and sequenced on an in-house MiSeq platform (lllumina, USA). Reads were trimmed, mapped, and processed as previously described with the Con-ARTIST TIS analysis pipeline (*13*). To identify synthetic interaction loci, we used WT as the “input” library and Δ*vca0040* as the “output” library during analysis.

#### RNAseq

For transcriptomic analyses, triplicate WT and Δ*vca0040* V. cholerae 37°C overnight cultures were back-diluted 1:100 into fresh LB medium and grown at 30°C for 8 hours with shaking. At 8 hours, samples were checked by microscopy to ensure onset of sphere formation in the mutant samples, at which point 2 mL of each culture was spun down (5 minutes at 5000 rcf at RT). RNA was extracted with Trizol reagent (Sigma-Aldrich, USA) per the manufacturer’s instructions. Isolated RNA was then DNase-treated and re-isolated with ethanol precipitation according to a standard protocol. RNA samples were quality checked with a Bioanalyzer to confirm RIN values > 6. Library preparation and sequencing was performed by the Microbial Genome Sequencing Center (Pittsburgh, USA). RNAseq analysis was performed largely as described (*43*). Reads were mapped 7 to the *V. cholerae* KW3 genome (NCBI assembly GCA_001318185.1) with Bowtie2 (Galaxy) and a read matrix was generated with featureCounts (*44*). Read matrices were then analyzed with the default DESeq2 pipeline in RStudio to identify differentially expressed genes. Data shrinkage was performed with ashr (*45*).

#### Multiplexed tandem mass tag (TMT) proteomics

For proteomic analyses, triplicate WT and Δ*secDF1* V. cholerae 37°C overnight cultures were grown as described for RNAseq. At 8 hours of growth, whole cell pellet (WCP) samples were prepared by centrifuging 1 mL of cells for 5 minutes at 5000 rcf at RT, washing once in fresh LB, and flash frozen and kept at −80°C. MS analysis was performed by the Thermo-Fisher Center for Multiplexed Proteomics (TCMP) at Harvard Medical School according to standard protocols. WCP samples were subjected to a total proteomics workflow with fractionation. Pelleted cells were first lysed in 8 M urea, 200 mM EPPS and 1% SDS with phosphatase and protease inhibitors. Then, samples were reduced with DTT and alkylated with iodoacetamide. Alkylated proteins were precipitated with methanol/chloroform and resuspended in 200 mM EPPS pH 8 and digested sequentially with 1:50 LysC and 1:100 trypsin. Peptides were labeled with TMT16 reagents, pooled, and fractionated by a basic reverse phase (bRP) protocol into 12 fractions. Fractions were then dried, cleaned on a C18-packed stage tip, and eluted into an MS sample vial for analysis. Samples were resuspended in 5% ACN/5% formic acid and analyzed by LC-MS3 on an Orbitrap Lumos mass spectrometer. Peptides were detected (MS1) and quantified (MS2) in the Orbitrap, and sequenced (MS2) in the ion trap. MS2 spectra were searched with the COMET algorithm against the *V. cholerae* KW3 proteome, its reversed complement, and known contaminants. Spectral matches were filtered to a 1% false discovery rate using the target-decoy strategy combined with linear discriminant analysis. Proteins were quantified from peptides with a summed signal to noise threshold of >150 and isolation specificity of >0.5.

#### Spontaneous suppressor isolation and sequencing

##### V. cholerae

To ensure independent isolation of spontaneous suppressors, the Δ*vca0040* strain was re-derived 11 times from different conjugation reactions. From each colony PCR- and cell shape defect-verified Δ*vca0040* clone, colonies with mutant morphology (small and completely blue) on LB+Sm/X-gal plates were re-streaked onto new LB+Sm/X-gal plates and grown for 36-48 hours at 37°C. This process was repeated once. From the tertiary plates, a single colony with WT morphology (large with a white halo) was re-streaked to confirm a stable suppressor phenotype. Suppressors were checked by microscopy of 30°C overnight cultures to confirm cell shape defect reversion and verified by colony PCR to confirm the absence of *vca0040* from their genome. Validated suppressors were grown overnight, and genomic DNA was extracted as described above. Library preparation and sequencing were either outsourced to the Microbial Genome Sequencing Center (Pittsburgh, USA) or performed in-house. For in-house whole genome sequencing, gDNA was tagmented and barcoded with the Nextera XT library preparation kit (Illumina, USA), quality checked by BioAnalyzer and sequenced on a MiSeq platform to at least 20-50x depth of the *V. cholerae* genome (∼4 Mbp). Genome assembly and variant identification was performed with CLC Genomics Workbench 12 (Qiagen, Germany). Variants were filtered against an assembled HaitiWT isolate re-sequenced with the same workflow. Mutations were assigned as suppressors if they were present in >90% of reads and did not occur in a known poorly mapping or highly varying region such as a tRNA locus. For the secondary screen in *secDF2-*Δ*vca0040* bacteria, the Δ*vca0040* deletion was re-derived three independent times in the Δ*secD2* or Δ*secF2* background. Suppressors were isolated identically to those in the parental Δ*vca0040* background.

##### S. aureus

Multiple 2 mL cultures of *S. aureus* HG003 Δ*SAOUHSC_00846* were grown in TSB overnight at 37°C. 190 µL from each independent overnight culture was then separately plated on TSA 100 mM Tris pH 9 plates (where the mutant is entirely inhibited for growth) and grown at 37°C for 24 hours. Suppressors were confirmed by re-streaking on TSA 100 mM Tris pH 9 plates. Validated suppressors were grown overnight, and genomic DNA was extracted as described above. Library preparation, sequencing, and variant identification was performed as described above for *V. cholerae*.

#### Infant rabbit *V. cholerae* infections and cecal fluid imaging

Infant rabbit oral infections with *V. cholerae* were performed as described previously (*10*). Briefly, 2–3-day-old New Zealand White rabbits (Charles River Laboratories) co-housed with their dams were orally gavaged with 10^9^ CFU *V. cholerae* in a 500 μL gavage volume. Inocula were prepared by centrifuging a late exponential phase culture (OD_600_ 0.6-0.8) *V. cholerae* and resuspending in 2.5% sodium bicarbonate. For varying the pH of the inocula, sodium bicarbonate was pH-adjusted to pH 9 or 7 with NaOH or HCl immediately before resuspension. Infected kits were returned to their dam and monitored for 16-20 hours post-infection, when they were sacrificed by isoflurane inhalation and intracardiac injection of 20 mEq potassium chloride. Small intestinal segments and the cecum were isolated by dissection, homogenized by bead beading in PBS, and dilutions were plated on appropriate agar plates for colony enumeration. Plates were counted after an overnight incubation at 30°C. During dissection of the cecum, a 28 G needle was used to extract crude cecal fluid (CF). A sample of crude CF was taken for CFU plating and imaging, and the remainder was centrifuged at 21000 rcf for 2 minutes. The pellet was discarded and supernatants were frozen at −20°C for downstream analyses. For CF incubation assays, overnight cultures of *V. cholerae* were back-diluted 1:100 in CF and grown for 4 hours at 30°C shaking before imaging.

#### Mouse *S. aureus* intravenous infection

*S. aureus* HG003 (WT) and KanR-marked Δ*SAOUHSC_00846* were grown overnight at 37°C. Cultures were mixed at a 1:1 ratio, combined 1:1 with 50% glycerol, and stored at frozen at −80°C in several aliquots. For infections, an aliquot was thawed and diluted in PBS to a density of ∼3×10^7^ CFU/ml. 100 μl was injected into the tail vein of 8–9-week-old female Swiss Webster mice. Mice were monitored daily and at days 2 and 5 post-infection, the heart, lungs, right kidney, liver, and spleen were collected. Organs were homogenized with stainless steel beads in PBS and plated on TSA and TSA + kanamycin (TSAK) and incubated overnight to enumerate total Δ*SAOUHSC_00846* CFU, respectively.

#### Animal use statement

All animal work in this study was performed in accordance with the NIH Guide on Use of and Care for Laboratory Animals and with the approval of the Brigham and Women’s Hospital IACUC (Protocol #2016N000334 for infant rabbits and #2016N000416 for mice).

#### Statistics

Statistical tests used and replicate information are indicated in the figure legends and relevant methods sections. Statistical analyses were performed in Prism (Graphpad, USA).

### Supplementary Text

#### Variable annotation of coding sequences in the reference *V. cholerae* genome

During our study, which mainly used a contemporary pandemic strain of *V. cholerae* (HaitiWT or KW3, GCA_001318185.1), we noted several annotation inconsistencies with genes ostensibly conserved in the reference shotgun assembly (GCA_000006745.1) of N16961 *V. cholerae*. We found that compared to newer assemblies of N16961 (GCA_003063785.1 and GCA_900205735.1), the original assembly has ∼50 more ORFs annotated as pseudogenes due to frameshifts. Although these ORFs are still annotated, these loci lack protein accessions and are not included in coding sequence tables. Three of these were directly related to our study: 1) *lacZ*, a widely-used reporter gene (VC2338), 2) *secF2* (VCA0692), and 3) *yghB* (VCA0534). *lacZ* is known to be functional in N16961 and has been used repeatedly as a reporter locus. Re-sequencing of our laboratory N16961 stock and examination of newer N16961 assemblies confirmed that *secF2* and *yghB* are in fact intact in this strain and lack the predicted frameshift. Notably, the “pseudogene” annotation of *secF2* led to its usage as a safe-harbor locus for genome editing in *V. cholerae* in a recent report (*46*), but caution should be used for this method. The newer, long-read hybrid assemblies of N16961 *V. cholerae* may circumvent these issues and improved sequence and annotation curation will benefit future studies (*47*).

#### Phylogenetic conservation of DUF368 and DedA proteins

The conservation patterns and domain structures of DUF368 and DedA proteins support the claims in our manuscript for several reasons. First, we note that out of >30,000 representative bacterial genomes, just 2.8% of them lack an annotated DUF368, DedA or BacA/UppP (PF02673) domain. This is likely also an overestimate of the number of species lacking any of these three domains due to sequence divergence and inadequate PFAM annotation, which is not the most sensitive form of conservation analysis. For example, upon manual curation, the obligate intracellular pathogen *Rickettsia rickettsii* (GCA_000018225.1) is in the triply-absent group, but is known to have PG and has an annotated DedA coding sequence, perhaps reflecting a PFAM annotation lag. Interestingly, however, *Mycoplasma* species, which do not have LPS or PG (and thus may not be dependent on C55-P translocation) appear to be genuinely triply-absent except for a select few members with a DUF368 only. Second, DUF368 proteins in archaea (identified by Annotree) appear to be heavily enriched in the Halobacteria, which are known alkaliphilic halophiles, a niche consistent with our findings in *V. cholerae* and *S. aureus* (both known halophiles), suggesting that the presence of DUF368-containing proteins correlates with niche diversification toward alkaline and/or sodium-rich contexts. We also note that DUF368 is common in other known bacterial halophiles in the genus *Salinibacter* and order Halanaerobiales. Finally, using domain structure visualization on PFAM (pfam.xfam.org), DUF368 (PF04018) and DedA (PF09335) proteins are occasionally found fused to PAP2 (PF01569) C55-PP phosphatase domains. Beyond reinforcing the idea that these two protein families are C55-P translocases, this suggests that, similar to what has been proposed for UppP (*4*), phosphatase activity is coupled to transport by select DUF368 and DedA proteins.

### Supplementary Figures

**Fig. S1.**
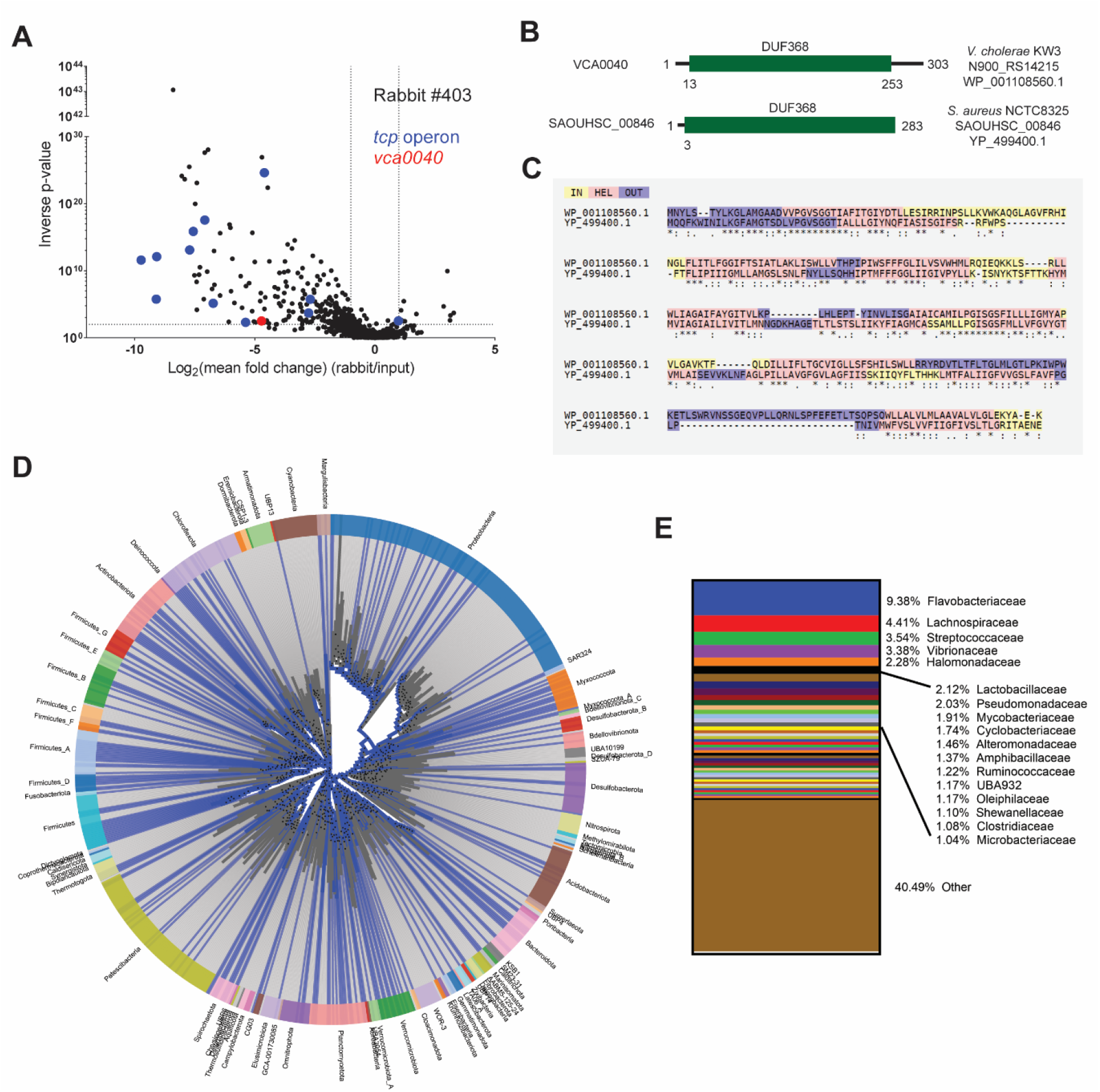
Identification and conservation of VCA0040. (A): Infant rabbit transposon-insertion sequencing data reproduced from Hubbard *et al.* (reference 10) identifying *vca0040* (red) as a *V. cholerae* intestinal colonization determinant. The *tcp* operon, known to be critical for colonization, is highlighted in blue. (B): Domain structure, locus tags, accession numbers, and alignment (C) of *V. cholerae* and *S. aureus* DUF368-containing proteins. Alignment was performed with T-COFFEE. IN: predicted cytosol-facing sequence. HEL: predicted transmembrane helix sequence. OUT: predicted external-facing sequence. (D and E): Phylogenetic conservation of DUF368 domains (PF04018). (D): Family-level conservation (blue lines) grouped by phylum (colored arcs). (E): Family-level breakdown of DUF368 conservation.

**Fig. S2.**
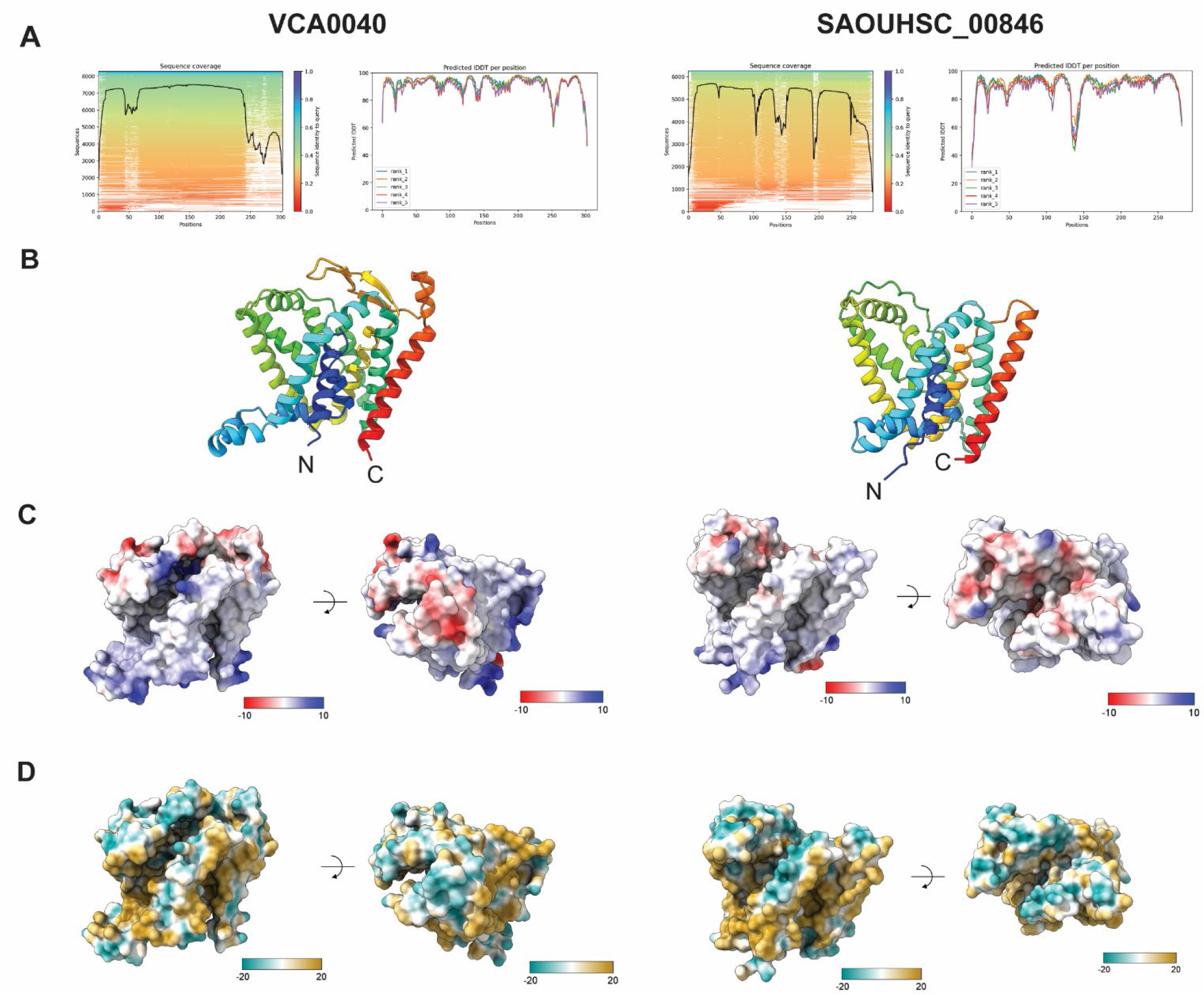
Structural modeling of DUF368-containing proteins. The *V. cholerae* (left) and *S. aureus* (right) DUF368-containing proteins were modeled using AlphaFold2 and visualized with ChimeraX. (A): MSA coverage and pLDDT per position for each protein from AlphaFold2. (B): Overall ribbon structures, colored from N (violet) to C (red) termini. (C and D): Electrostatic surface potential (C) and hydrophobicity (D) maps generated using default ChimeraX settings.

**Fig. S3.**
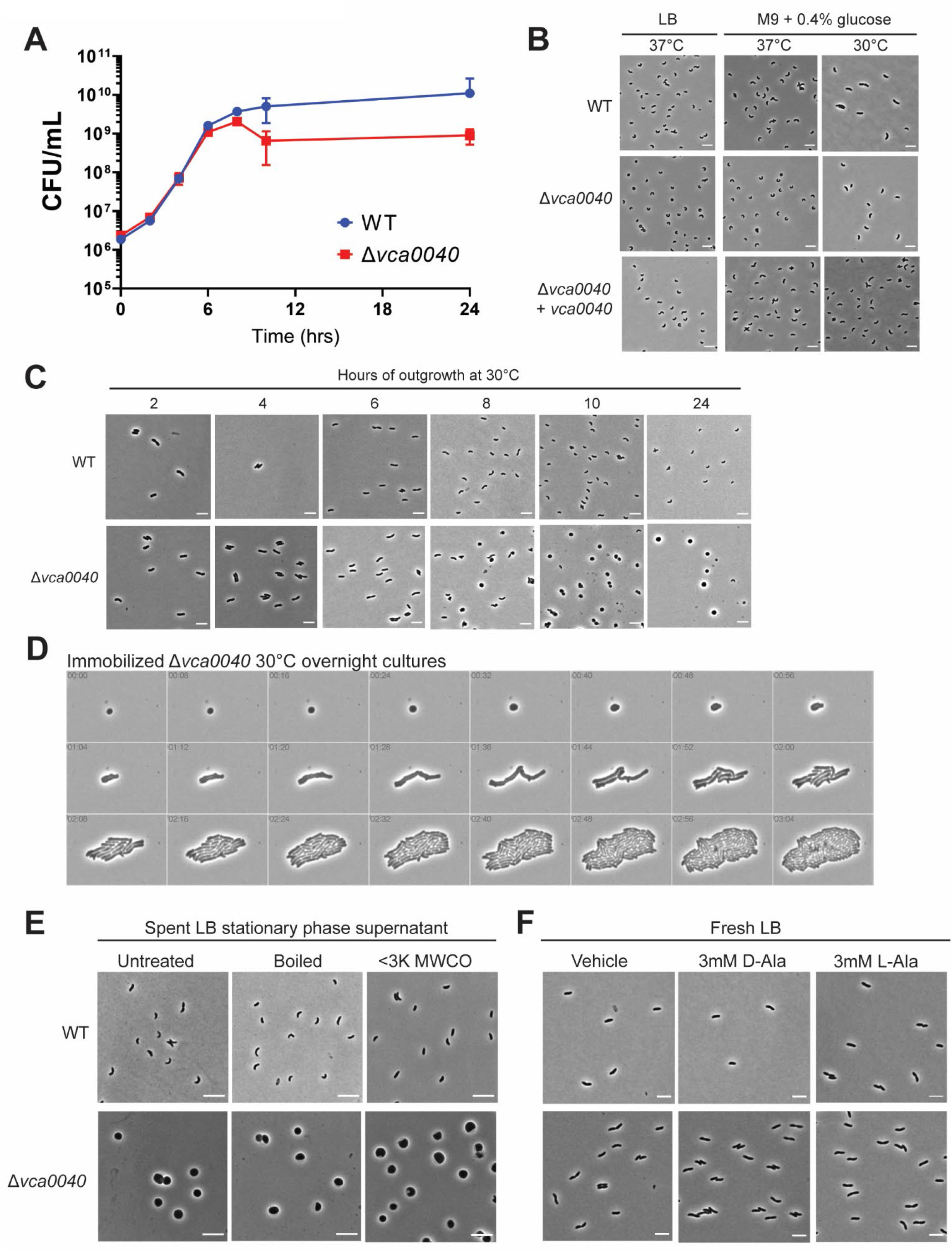
Growth of a *V. cholerae* strain lacking *vca0040.* (A): CFU plating time course of WT and Δ*vca0040 V. cholerae*. Arrow indicates onset of stationary phase. (B): Phase-contrast imaging of WT, Δ*vca0040* and Δ*vca0040+vca0040 V. cholerae* in overnight cultures in the indicated conditions. (C): Phase-contrast imaging time course of the indicated strains grown in LB at 30°C for the indicated amount of time. (D): Time lapse imaging of Δ*vca0040* cells immobilized on an agarose pad from a 30°C overnight culture. (E): Imaging of *V. cholerae* treated with normal, boiled (10 minutes) or filtered (<3K MW cutoff) cell-free spent supernatant for 4 hours at 30°C. (F): Imaging of *V. cholerae* treated with vehicle (DMSO) or 3mM of D- or L-Ala in fresh LB for 4 hours at 30°C.

**Fig. S4.**
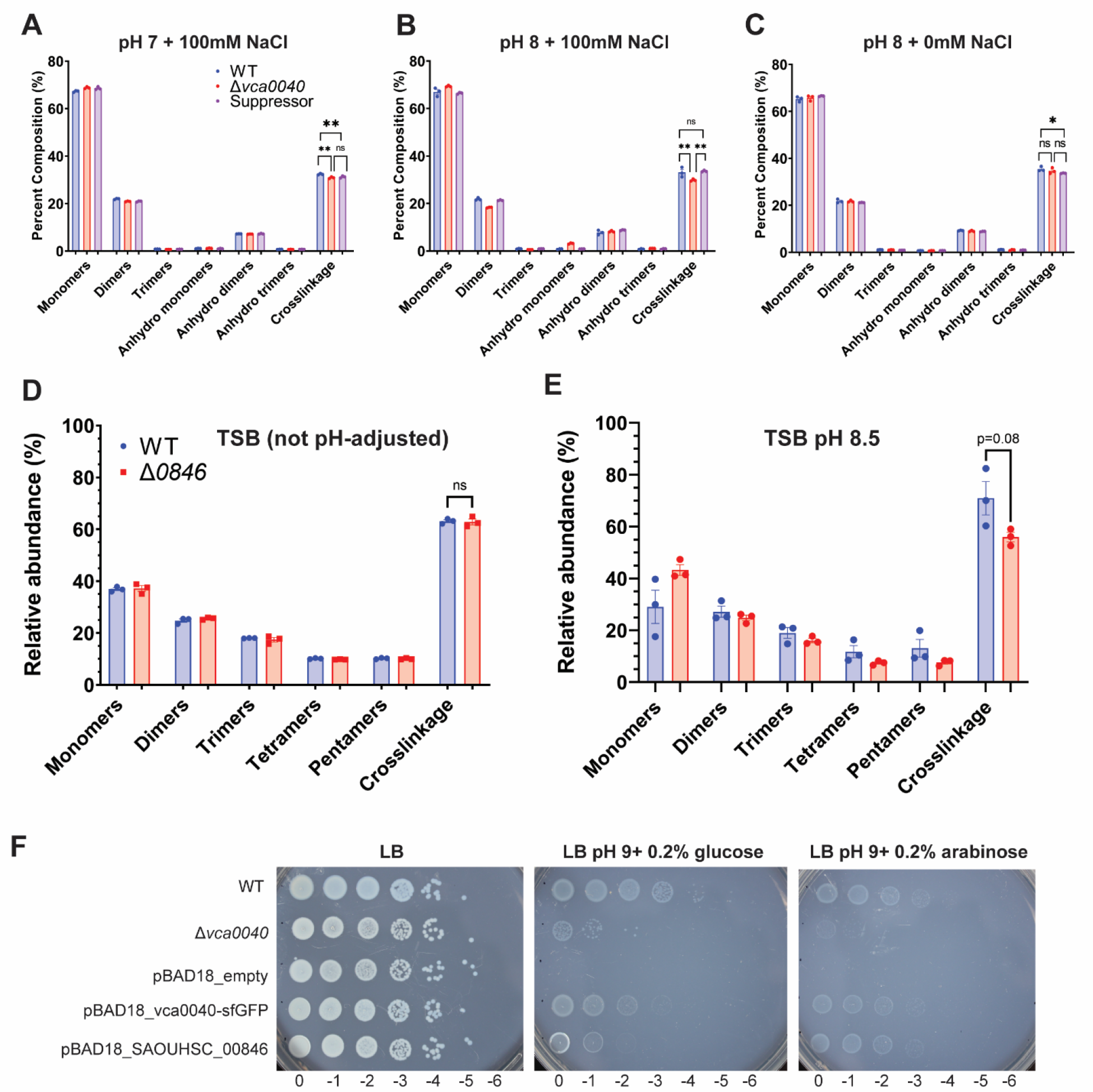
Analysis of PG composition and crosslinking in *V. cholerae* and *S. aureus.* (A-C): Composition and crosslinking analysis in the indicated *V. cholerae* strains at the indicated conditions. Suppressor: Δ*vca0040/ΔsecDF1 V. cholerae.* (D and E): Same as A-C for *S. aureus* grown in the indicated conditions. *p<0.05, **p<0.001 (unpaired student’s two-tailed t-test). ns, not significant. (F): Rescue of *vca0040 V. cholerae* alkaline defect by plasmid-based overexpression of *vca0040* or *SAOUHSC_00846*). The pBAD18_*vca0040-*sfGFP vector contains a C-terminal fusion with sfGFP for unrelated imaging studies. LB was buffered with 100mM Tris and adjusted with NaOH.

**Fig. S5.**
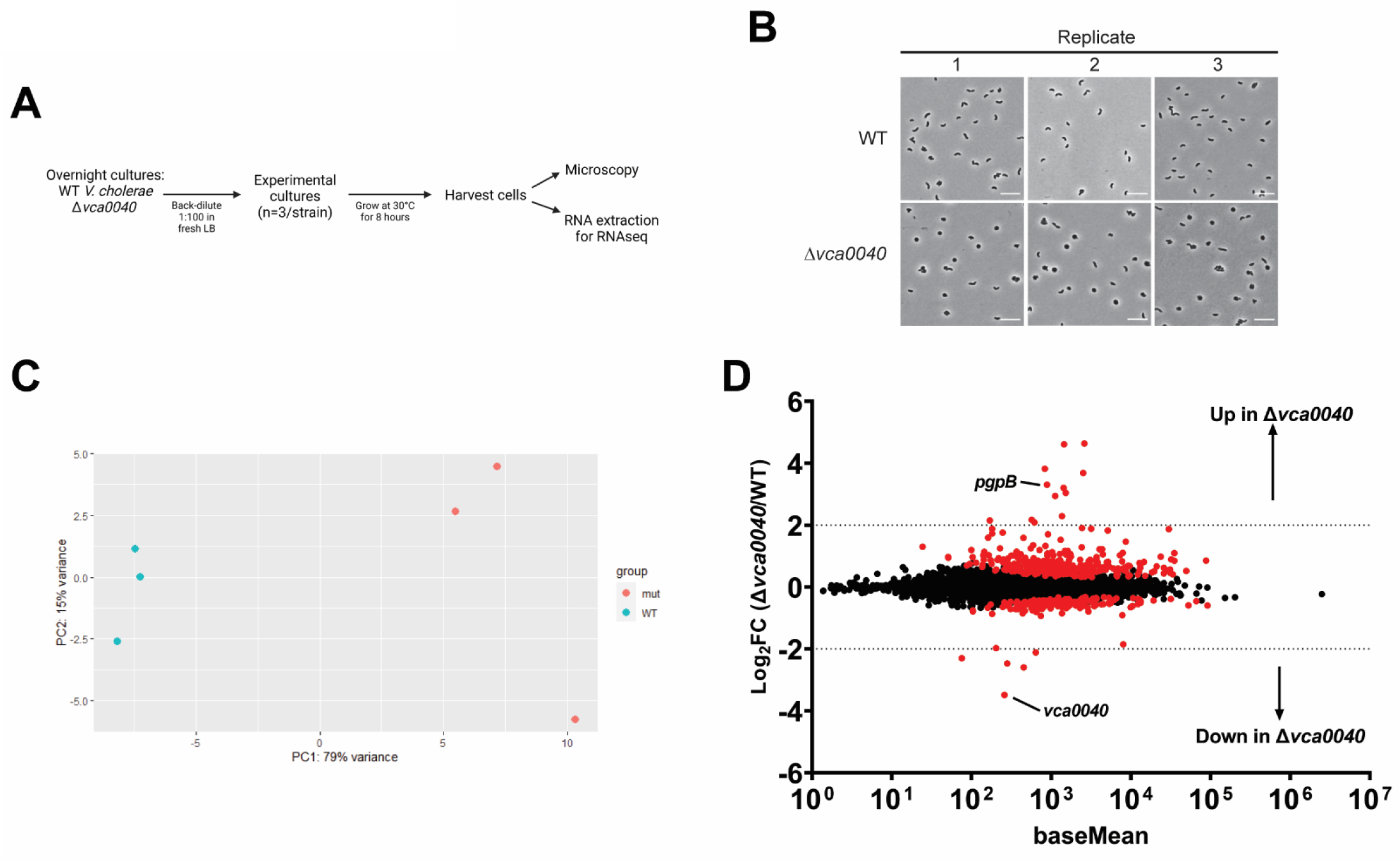
RNAseq of Δ*vca0040 V. cholerae* during sphere formation. (A – D): RNAseq of Δ*vca0040 V. cholerae,* with schematic (A), confirmation of sphere formation via imaging at sample collection (B), principal component analysis of RNAseq data (C), and M-A plot of analyzed sequences (D). An arbitrary fold change (FC) cutoff of 4 was implemented during RNAseq analysis, and genes with FC values beneath the p-value threshold of 0.05 are highlighted in red. *pgpB* and *vca0040* are highlighted. Data for this panel are available in Supplementary Table S5.

**Fig. S6.**
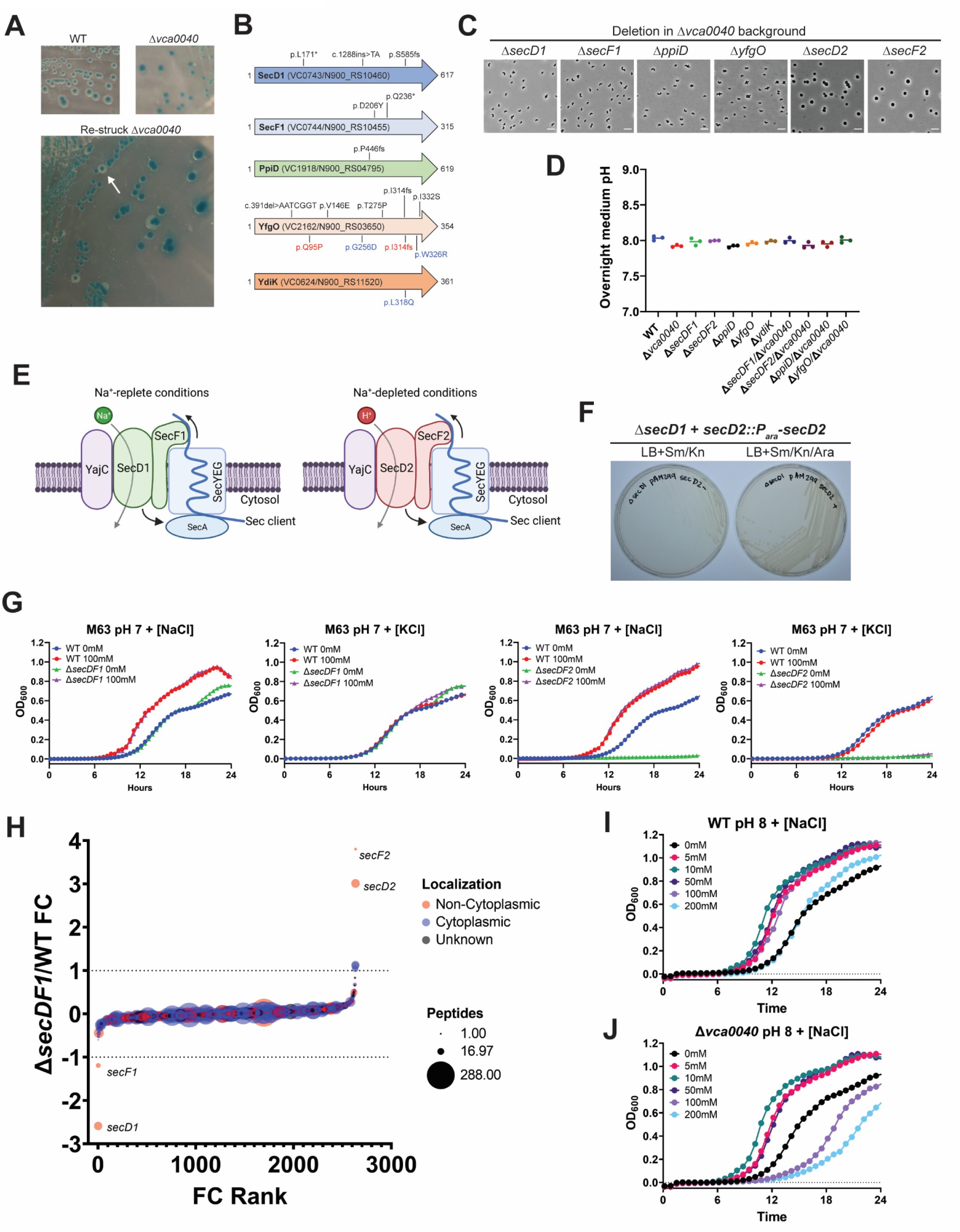
Identification and analysis of *vca0040* suppressor loci. (A): Representative colony morphologies of WT and Δ*vca0040 V. cholerae* on LB+X-gal plates. The white arrow indicates a suppressor colony. (B): Mutations identified in spontaneous suppressors of Δ*vca0040 V. cholerae.* SecD1, SecF1 and PpiD are part of the *V. cholerae* protein secretion machinery. The colored text indicates suppressors harvested in a secondary screen, where either a Δ*secD2* (blue) or Δ*secF2* (red) Δ*vca0040* strain was selected to bias suppressor mutations away from *secDF1* due to synthetic lethality. Asterisk: stop codon. (C): Phase-contrast imaging of 30°C LB overnight cultures of the indicated strains in the Δ*vca0040* background. (D) Overnight 30°C LB culture pH of the indicated strains. (E): SecDF1 and SecDF2 functions in *V. cholerae.* Curved arrows represent possible functions of SecDF: coupling ion import to the stimulation of SecA ATPase activity (bottom arrow) or physical pulling of a nascent Sec substrate into the periplasm (top arrow). (F): Synthetic lethality of *secD2* in Δ*secD1 V. cholerae* using the pAM299 system. (G): Growth of the indicated strains in M63 minimal medium with added NaCl or KCl. (H): Waterfall plot comparing Δ*secDF1* whole cell proteome to that of WT *V. cholerae*. (I and J): Growth of WT (I) or Δ*vca0040* (J) *V. cholerae* in M63 pH 8 with a range of NaCl concentrations. Note that the Δ*vca0040* strain can only tolerate up to 10 mM NaCl before exhibiting a growth defect.

**Fig. S7.**
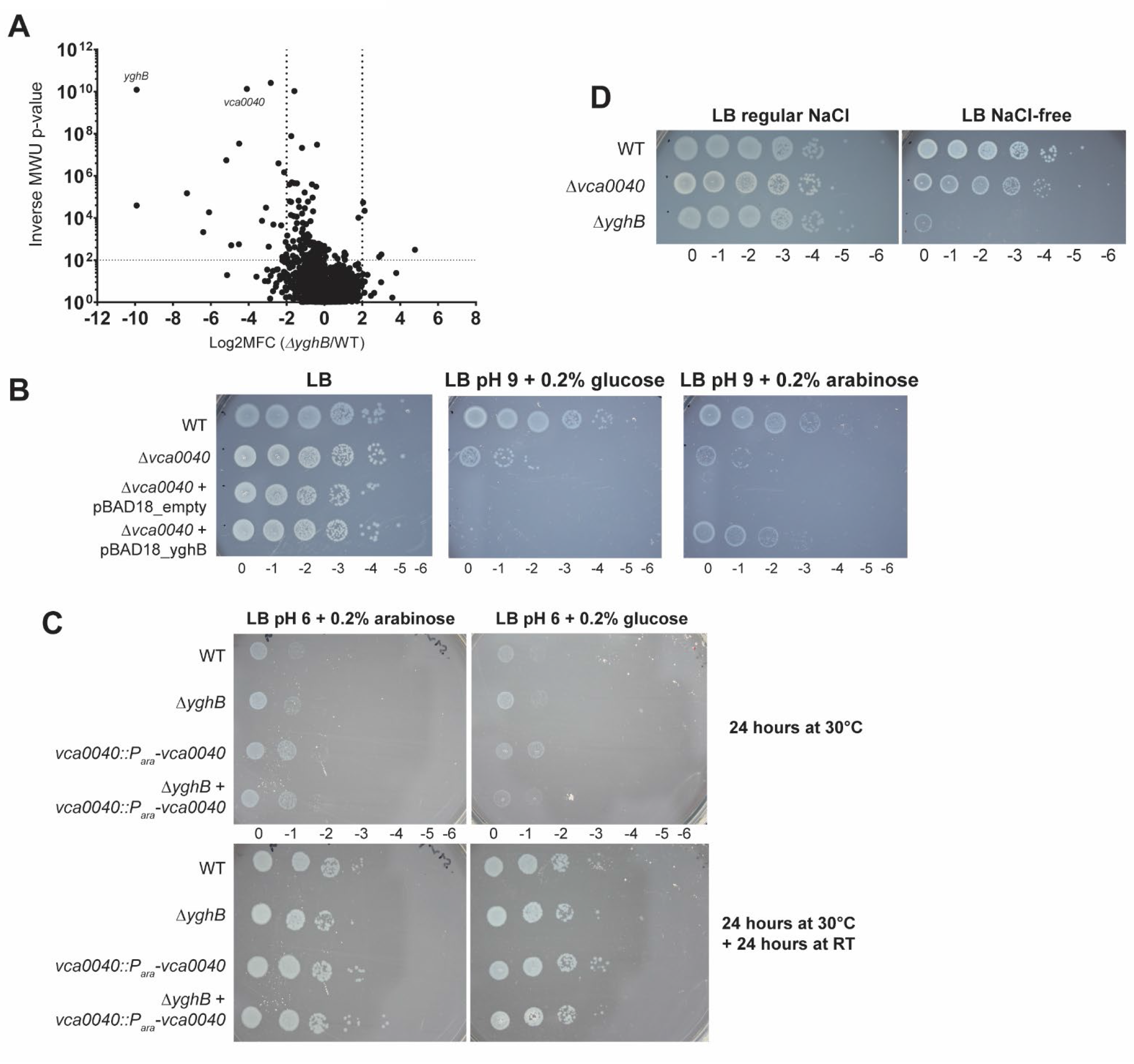
Genetic interactions between DedA and DUF368 proteins. (A): Synthetic transposon-insertion screen in Δ*yghB V. cholerae.* (B): Growth of Δ*vca0040 V. cholerae* expressing vectors with arabinose-inducible genes on regular LB (left), LB pH 9 (middle) and LB pH 9 + arabinose (right) plates. (C): Growth of *V. cholerae* lacking one or both of *vca0040* and/or *yghB* on acidic LB plates. LB was pH-adjusted with HCl and buffered with 100 mM MES. (D): Growth of the indicated *V. cholerae* strains on LB with regular [NaCl] (∼170 mM) and no added NaCl.

**Fig. S8.**
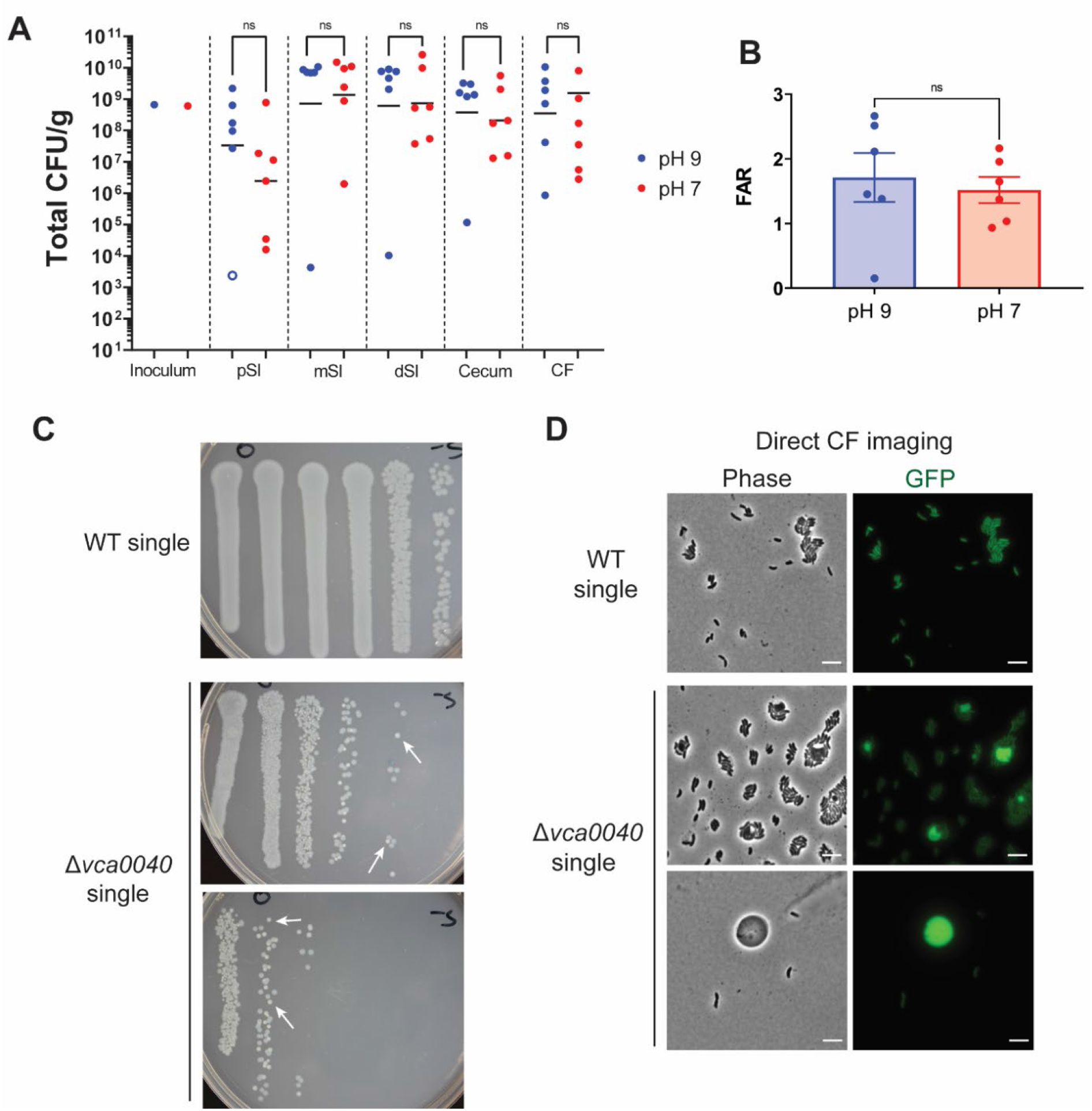
Supplemental data for infant rabbit infections. (A): Total CFU in mixed infections with differing inoculum pH. (B): Fluid accumulation ratio (FAR) in mixed infections with differing inoculum pH. (C): Representative small intestinal CFU plates showing transmission of WT *V. cholerae* (greyish translucent colonies) from WT-infected rabbits to Δ*vca0040*-infected rabbits in the same litter. Note that Δ*vca0040* colonies (white arrows) are lighter and smaller than WT colonies, making them visually distinguishable. (D): Direct imaging of GFP+ *V. cholerae* in single infection CF samples. Large GFP-bright spheres are likely Δ*vca0040* cells in the presence of transmitted, rod-shaped WT cells. ns, not significant (Mann-Whitney U test). Scale bars, 5μm.

**Fig. S9.**
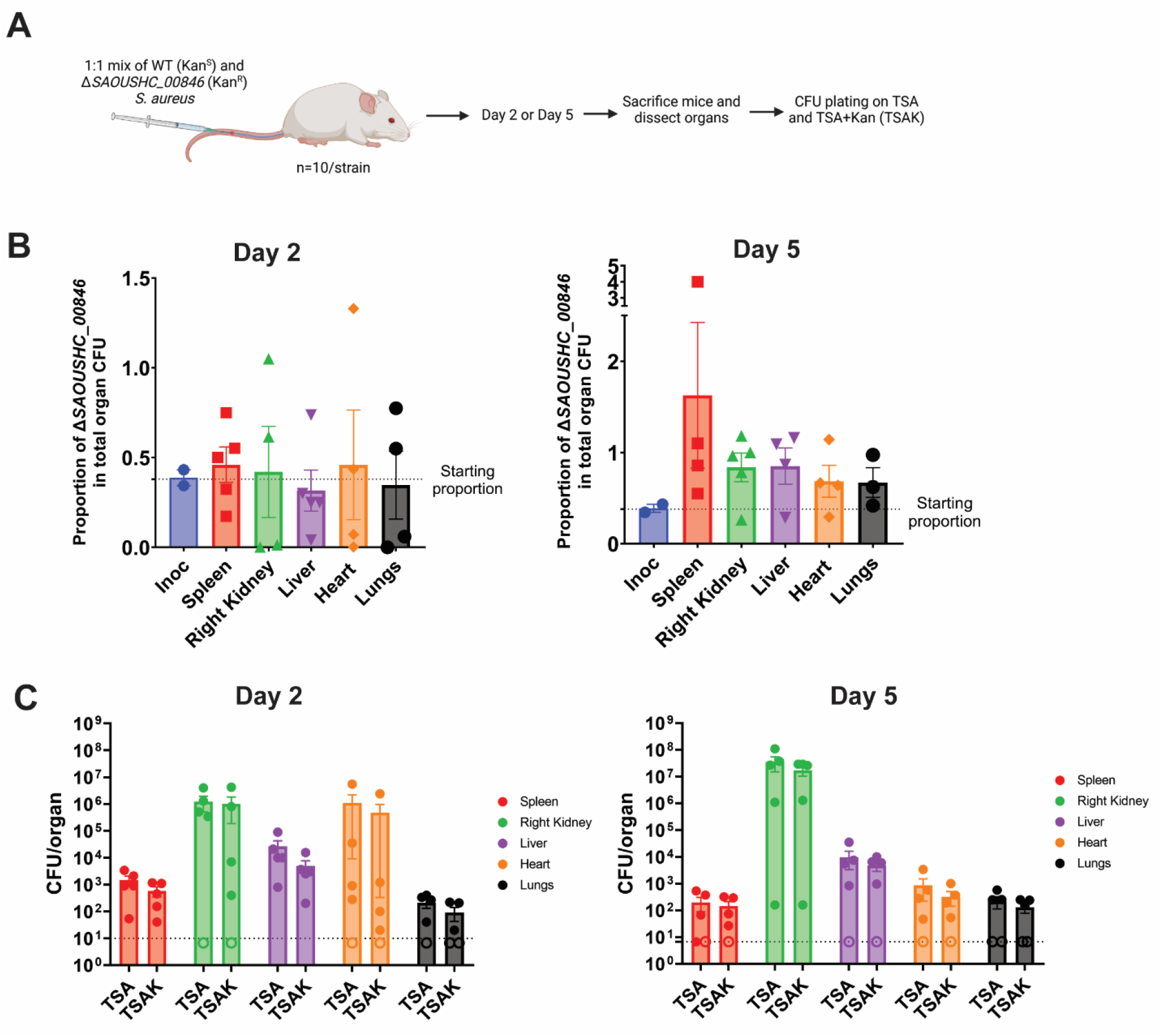
Intravenous (IV) infections of mice with *S. aureus.* (A): Schematic of IV infections and sample harvesting workflow. TSA counts indicate total *S. aureus* CFU burden, whereas TSAK counts indicate Δ*SAOUHSC_00846* burden. (B): Ratio of Δ*SAOUHSC_00846* (KanR) colony-forming units (CFU) to total *S. aureus* CFU at Day 2 (left) and Day 5 (right) post-infection. (C): Raw CFU counts from TSA and TSA + kanamycin (TSAK) plates. In the absence of a competitive defect, the TSAK counts should be 50% of those on the TSA plates (and thus are very close together on the log_10_-plotted graph).

**Fig. S10.**
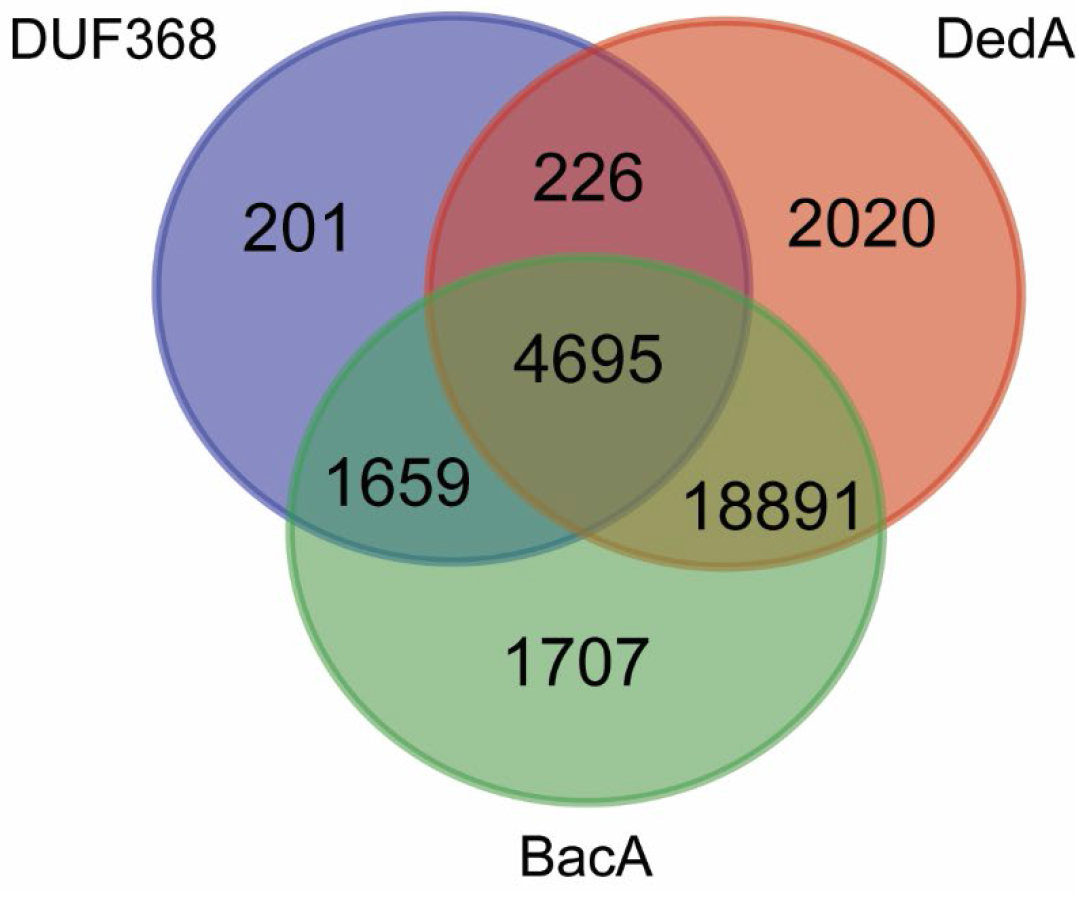
Distribution of putative C55-P translocases in bacteria. Venn diagram of Annotree species-level outputs for PFAM queries PF04018 (DUF368), PF09335 (DedA), and PF02673 (BacA/UppP). Out of 30,238 bacterial species in the Annotree database, 29,399 are predicted to contain at least one of the three proteins. A list of species in each Venn diagram segment is **listed in table S9**.

**Fig. S11.**
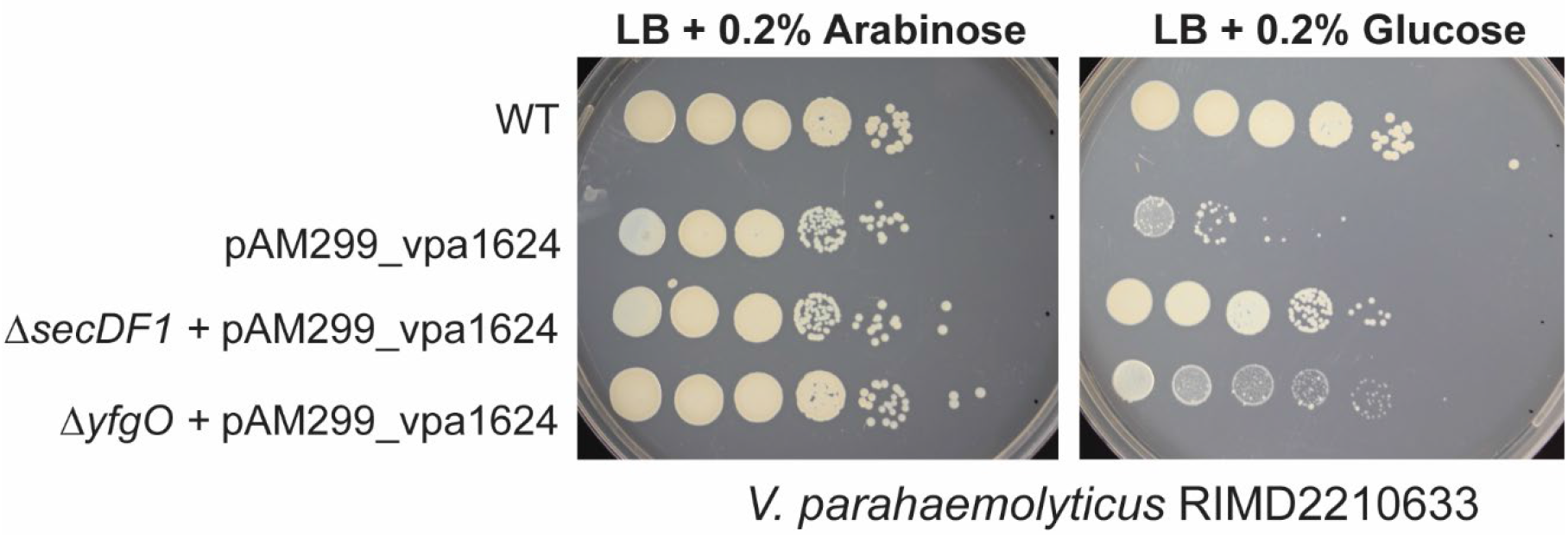
Essentiality and genetic suppression of a DUF368 protein in *V. parahaemolyticus.* The *V. parahaemolyticus* DUF368 protein, VPA1624, was depleted with the pAM299 system and is clearly required for growth even on neutral LB plates. VPA1624 apparently genetically interacts with the *V. parahaemolyticus secDF1* and *yfgO* loci, as overexpression of these suppressors originally identified in *V. cholerae* rescues the VPA1624-depleted strain.

### Supplementary Tables

All supplementary tables are included as additional spreadsheets with the submission.

**Table S1. Bacterial and archaeal clades with PF04018-containing proteins.**

Annotree was used to plot and identify species with an annotated DUF368 domain. Each tab of the spreadsheet is a different level of bacterial classification (Phylum>Order>Family>Genus>Species). Archaeal clades are combined on the tab “Archaea”.

**Table S2. Homologues of VCA0040 in bacteria.**

HMMER was used to identify homologous sequences to VCA0040 in the Uniprot database. Each species is listed once.

**Table S3. MICS for *V. cholerae* WT and Δ*vca0040*.**

**Table S4. MIC data for *S. aureus* WT and Δ*SAOUHSC_00846* and *E. coli* WT and *lptd4213*.**

**Table S5. RNAseq of Δ*vca0040 V. cholerae* during sphere formation.**

For the top 40 up-regulated genes in Δ*vca0040*, manual curation of subcellular localization was performed with SignalP and pSortb.

**Table S6. Multiplexed comparative proteomics of whole-cell WT and Δ*secDF1 V. cholerae*.**

Fold change was calculated by dividing the average normalized relative abundance of proteins in Δ*secDF1* by that of WT.

**Table S7. Synthetic transposon-insertion screening in Δ*vca0040 V. cholerae*.**

Note that intergenic regions (IG_) and regions with <5 informative sites (possible transposon insertion sites) were excluded from analysis. Hits on the second tab were thresholded at an inverse p-value >100 before sorting by mean log_2_ fold change.

**Table S8. Synthetic transposon-insertion screening in Δ*yghB V. cholerae*.**

Note that intergenic regions (IG_) and regions with <5 informative sites (possible transposon insertion sites) were excluded from analysis. Hits were thresholded at an inverse p-value >100 and log_2_FC less or greater than 1 (2-fold change).

**Table S9. Conservation of PF04018, PF09335, and PF02763 in bacterial species.**

Spreadsheet was generated by Annotree queries for all possible combinations of the three protein families. The “Master” sheet lists all species-level organisms with the indicated combination of domains. The “Model Species” sheet lists selected microbes of particular interest and can be used to look up any given species in the reference table. Note that species of interest should be manually confirmed for presence or absence by BLAST to guard against misannotation or non-annotation issues.

**Table S10. Strains and vectors and genomic information used in this study.**

**Table S11. BLAST dictionary for assignment of putative VC names to HaitiWT *V. cholerae* loci.**

All coding sequences from HaitiWT were used as queries in a batch BLAST of the N16961 *V. cholerae* proteome. The top hit by e-value was selected and VC name automatically assigned. However, as batch BLAST outputs a hit regardless of confidence, manual curation is still required for specific loci when using this dictionary.

